# Exploration of endogenous miRNA-200b/c activity and regulation through a functional dual fluorescence reporter

**DOI:** 10.1101/2020.07.19.210997

**Authors:** Paradesi Gollavilli, Beatrice Parma, Aarif Siddiqui, Hai Yang, Vignesh Ramesh, Francesca Napoli, Annemarie Schwab, Ramakrishnan Natesan, Irfan Ahmed Asangani, Thomas Brabletz, Christian Pilarsky, Paolo Ceppi

## Abstract

Since their discovery, microRNAs (miRNA)s have been widely studied in almost every aspect of biology and medicine, leading to the identification of important gene regulation circuits and cellular mechanisms. However, investigations are generally focused on the analysis of their downstream targets and biological functions in overexpression and knockdown approaches, while miRNAs endogenous levels and activity remain poorly understood. Here, we used the cellular plasticity-regulating process of epithelial-to-mesenchymal transition (EMT) as a model to show the efficacy of a fluorescent sensor to separate cells with distinct EMT signatures, based on miR-200b/c activity. The system was further combined with a CRISPR-Cas9 screening platform to unbiasedly identify miR-200b/c upstream regulating genes. The sensor allows to infer miRNAs fundamental biological properties, as profiling of sorted cells indicated miR-200b/c as a molecular switch between EMT differentiation and proliferation, and suggested a role for metabolic enzymes in miR-200/EMT regulation. Analysis of miRNAs’ endogenous levels and activity could lead to a better understanding of their biological role in physiology and disease.

## INTRODUCTION

MicroRNAs (miRNAs) are short, highly conserved, RNAs that negatively regulate the expression of their target genes. Depending on the binding efficiency, miRNAs can either cause degradation of the target mRNA or inhibit translation resulting in effective down regulation (1). MiRNAs directly or indirectly regulate many cellular processes like inflammation, cell cycle, stress response, apoptosis, senescence, aging and migration (2,3), and have crucial functions in the development of different cancers. MiRNA can work as tumor suppressors or oncomiRs (4–6), and have been linked with all hallmarks of cancer, like proliferation, cell survival, evasion of apoptosis, angiogenesis, migration and invasion (6–11). Cancer-specific miRNA signatures characterize their malignant state defining key clinicopathological features like grade, stage, aggressiveness, vascular invasion, proliferation index and others (12).

Epithelial to mesenchymal transition (EMT) is an essential embryonic development program, which many cancers hijack and turn into their advantage to increase the migratory and invasive capacities of cancer cells (13). EMT can be induced by several signalling pathways, including TGF beta, Notch, and Wnt pathways (14,15). All these pathways culminate into reducing cell adhesion molecules, like E-cadherin. EMT is enforced by specialized transcription factor families SNAIL, basic helix loop helix (bHLH), TWIST and ZEB (16,17), which can work as E-cadherin repressors and promote the expression of EMT effector genes. MiRNAs are known to affect the EMT process by regulating the expression of one or more EMT-inducing transcription factors (16,18). For example, miR-200 inhibits the transcription factors of the ZEB family, in a reciprocal feedback loop (19) and miR-34 is a well-known inhibitor of EMT which acts by targeting SNAIL (20). The equilibrium between EMT-suppressing miRNAs and key EMT transcription factors determines the epithelial or mesenchymal state of the cells and their cellular plasticity, see **Figure 1A**.

**Figure 1.**
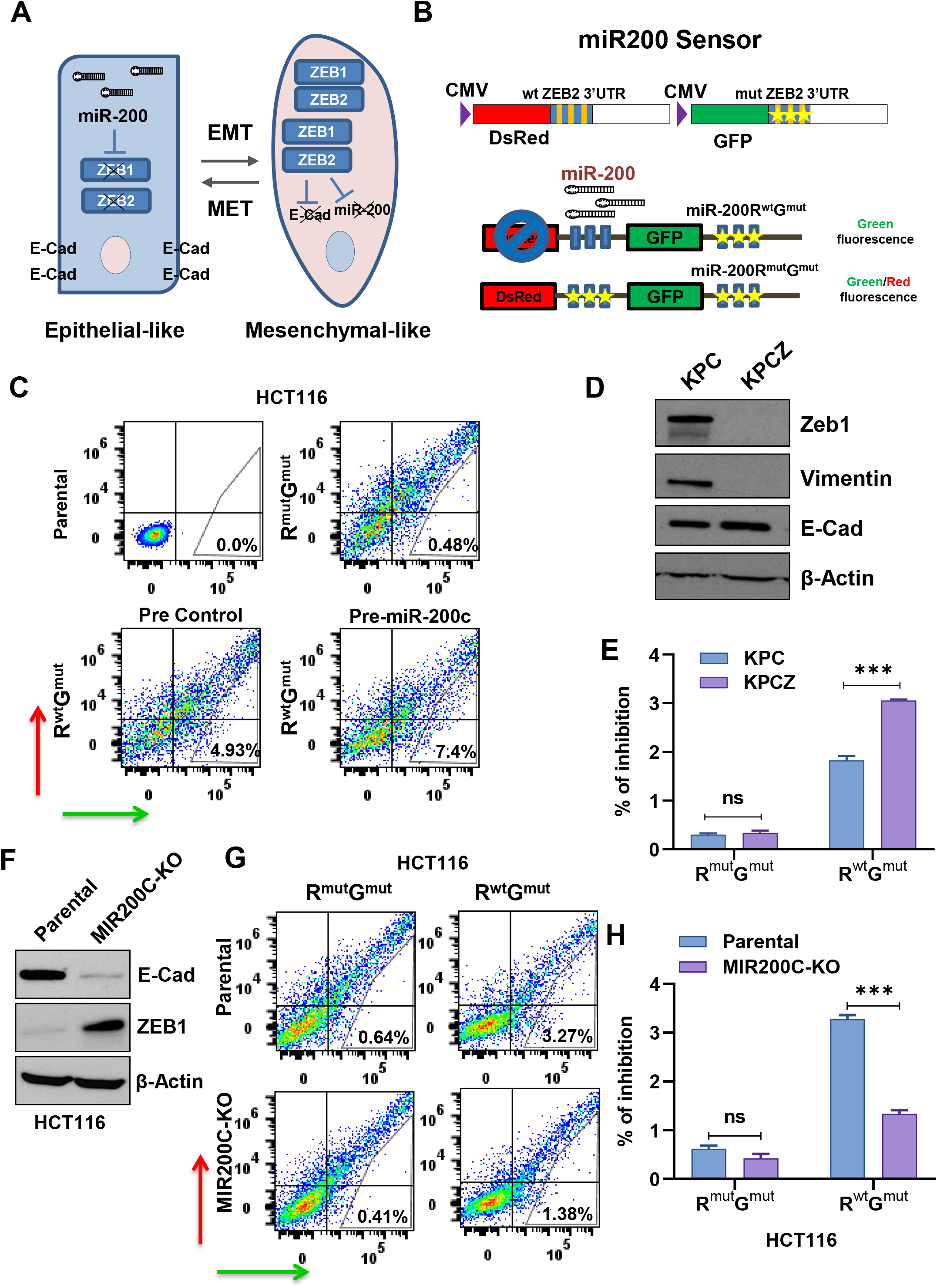
The miRNA sensor plasmid can be transiently transfected in cells to detect miR-200b/c levels to distinguish differential EMT states. A) Schematics showing the role of miR-200 in the regulation of epithelial to mesenchymal transition (EMT) and the feedback loop with ZEB1/2 E-Cadherin repressors. B) Schematic representation of the miR-200b/c sensor. R^mut^G^mut^ and R^wt^G^mut^. wt=wild-type, mut=mutated, ZEB2 3’UTR=3’ untranslated region of ZEB2 gene. C) FACS plots showing the fluorescence intensity of HCT116 cells with FITC-A (Green) and PE-A (Red) channels after transfection with sensor plasmid in the presence of either pre-control or pre-miR-200c at 100nM concentration. Indicated are the % of gated cells (% of inhibition). D) Western blots showing abrogation of Zeb1 and Vimentin and upregulation of E-Cadherin in KPCZ compared to KPC cells. E) Bar graphs showing % of inhibition with miR-200b/c sensor in KPC and KPCZ cells. F) Western blot quantification of ZEB1 and E-cadherin protein expression of HCT116 cells with miR-200c knock out (MIR200C-KO), compared to control cells. β-Actin was used as loading control. G) FACS plots showing the transfection of sensor plasmids in MIR200C-KO HCT116 cells or in parental control cells. H) Bar graphs showing the % of inhibition of FACS analysis done in (G). In E p-values are from two-way ANOVA. In H p-values are from students *t*-test. *<0.05, **<0.01, ***<0.001, ****<0.0001

MiRNAs are usually functionally investigated by overexpressing either miRNA mimics or inhibitory knocking-down sequences, and the downstream effects are measured by biochemical identification of target genes and by the analysis of cellular phenotypes. However, high-throughput functional assessment indicates that the majority of miRNAs expressed in cells do not show detectable targeting activity (21), suggesting the need for robust miRNA functional reporters to better investigate their physiological role. Such tools could also allow unbiased identification of the regulatory network controlling each miRNA, with several important implications. The use of miRNA sensors has been recently shown as an effective method to detect miRNA activity in living cells (21–24), and in limited cases used to isolate cells with distinct biological properties (25). The systems so far proposed are based on artificial 3’UTRs containing multiple repeated perfect complementarity miRNA sequences or binding sites, instead of naturally occurring 3’UTRs, and generally lack target binding mutant controls. Such controls are, essential in miRNA target identification studies, as miRNA targeting can only be concluded by the loss of regulation in constructs with mutated miRNA target sites (26).

To develop a robust and controlled plasmid-based fluorescent miRNA sensor, we cloned a 3’UTR fragment of the miR-200 target ZEB2 (17), containing 3 strong miR-200 seed matches fused to DsRed (R) fluorescent protein, driven by a CMV promoter. As a non-targeting control, in the same plasmid we fused GFP (G) to the ZEB2 3’UTR carrying mutated seed matches, driven by an identical CMV promoter and named the R^wt^G^mut^ vector. Once expressed in living cells, this plasmid is designed to respond to miR-200 levels altering the intensity of the red fluorescence, while green fluorescence serves as transfection control. A double mutant non-binding control (R^mut^G^mut^) was also generated, **Figure 1B**. The R^wt^G^mut^ plasmid was found to have high specificity for the miR-200 b/c cluster, which differs from the miR-200a/141 cluster by one nucleotide only in the seed match region (27). In this study we report that EMT biology can be explored in living cells by the miR-200b/c sensor plasmid. These findings could be exported to virtually all biologically-relevant miRNAs in different cellular models, with several important applications for miRNA research.

## MATERIAL AND METHODS

### Cell culture and Chemicals

HCT116 (ATCC) cells were cultured in McCoy’s 5A, RT112 and COLO205 (ATCC) cells were cultured in RPMI1640, and HEK 293 (ATCC) were cultured in DMEM media. Media were supplemented with 10% FBS (Sigma), 1% Pencillin/Streptomycin (Sigma) and 1% L-glutamine (Sigma). Cells were cultured at 37°C and 5% CO_2_ in a humidified incubator. Cells were STR authenticated, used between 3 and 15 passages, and tested for mycoplasma regularly (detection kit from Invivogen). HCT116-Cas9 cell line was generated by lentiviral transduction with lentiCas9-Blast (Addgene # 52962) and selection with 4 μg/mL of blasticidin (Sigma) for 3 days. KPC (Pdx1-cre;Kras^LSL.G12D/+^;Tp53^LSL.R172H/+^) and KPCZ (Pdx1-cre;Kras^LSL.G12D^/+;Tp53^LSL.R172H/+^; ZEB^fl/fl^) cells are mouse pancreatic cancer cells without and with ZEB1 knockout, as previously described (28), and were cultured in DMEM. TNF-alpha was purchased from Gibco (Life technologies). Cdk inhibitor (CGP 60474) was purchased from Tocris (Biotechne).

### Transfection

The sensor plasmids were generated cloning on the pCDHCMV-MCS-EF1-Puro vector from BioCat as previously described (27).Transfection of sensor plasmids were done using polyethyleneimine (PEI) mixing plasmid and PEI in 1:3 ratio in 0.9% NaCl. After 20 minutes of incubation, the mixtures was added drop wise to the cells. Co-transfection of miRNA mimics (Ambion) and sensor plasmids was done with lipofectamine2000 (Life technologies). pCDNA3.1(+)-Hygro plasmids and empty vector control were purchased from Genscript and transfected with PEI. pCDNA3.1(+)-Hygro-transfected cells were selected and grown as stable cultures adding 250ug/ml hygromycin in the culture media.

### FACS analysis

HCT116 cells were seeded in 6 well plates at a density of 0.5 million per well. The next day cells were transfected with 1.5μg of either R^mut^G^mut^ or R^wt^G^mut^ plasmids. After 48 hours cells were trypsinized, washed and resuspended in FACS buffer (5 mM EDTA and 2% FBS/PBS). Samples were run on Cytoflex FACS machine (Beckman). FACS data were analysed using FlowJo software v10.6.

### Western Blot

Cells were lysed in RIPA buffer and protein concentration was measured using BCA kit (Thermo Fisher). Protein lysates were resolved on 8-10% gels and transferred to PVDF membrane. Membranes were blocked in 5% milk in TBST for 1 hour and incubated with the primary antibodies overnight. Membranes were then washed with TBST and incubated with secondary antibodies for 1 hour. The membranes were developed with ECL reagent (Thermo Fisher) on to X-ray films using the chemiluminescence imager, AGFA CP100. Antibodies used are, E-cadherin, Vimentin, β-Actin (HRP conjugated) from Cell Signalling Technologies, ZEB1 from Sigma-Aldrich, H6PD and GNPDA1 from Thermo Fisher, HSD17B1 from Abcam. Cas9 antibody (bD-20) was purchased from Santa Cruz Biotechnology. Horseradish Peroxidase (HRP)-conjugated secondary antibodies were from Southern Biotech.

### Proliferation assay

Cells were seeded in a 96 well plates at very low density (5-10% confluency). Proliferation was measured by IncuCyte ZOOM live cell imaging system (Essen BioScience) scanning every 2-4 hours. The phase contrast images acquired by IncuCyte ZOOM were used to train the ZOOM software to develop processing definition to mask the cells. The mask was further utilized to identify the cells in the images and determined the surface area at each scanned time point. The final output was represented as confluency percentage which was indicative of proliferation.

### Immunofluorescence

Immunofluorescence staining of cells grown on cover slips was performed as follows: HCT116 cells were grown on a glass cover slip until they were about 90% confluent. Cells were washed with PBS and fixed with 70% pre-chilled ethanol for 20 minutes at room temperature. Cells were blocked in 3% BSA in PBS for an hour after two PBS washes, followed by overnight incubation in 250th dilution of ZEB1 or E-cadherin (same Abs used for western blotting) in the blocking buffer. After the incubation, cells were washed with PBS and incubated with 250th dilution of secondary mouse or rabbit antibody in blocking medium for an hour. Cells were washed thrice with PBS and the cover slip was mounted on the glass slide with histology mounting medium (Fluoroshield with DAPI, Sigma). Slides were visualized and photographed using Leica DM5500B fluorescence microscope and merged using Leica Application Suite-X software.

### Luciferase assay

Luciferase assay for NF-κB was done using NF-κB reporter kit (BPS Bioscience) according to manufacturer’s instructions. Briefly, 25,000 HCT116 cells were seeded in each well of 96 well plate. Next day 1 μl of reporter plasmid was transfected using Lipofectamine2000 (Life Technologies). After 48 h the cells were lysed and luminescence was measured with Dual-Glo Luciferase assay system (Promega) according to the manufacturer’s instructions. Both Firefly and Renilla luminescence was measured and Firefly/Renilla ratio is taken as NF-κB activity.

### qPCR

Total RNA was extracted using miRNeasy kit (Qiagen) and 45ng of RNA was retro-transcribed using TaqMan MicroRNA reverse transcription kit (Applied Biosystems) with the respective primers of miRNAs from TaqMan-microRNA assays (Applied Biosystems). qPCR was performed using TaqMan probes for the respective miRNAs and Universal Mastermix (Applied Biosystems). RNU6B was used as internal control. For mRNAs, random hexamers primers and Tetro cDNA synthesis kit (Bioline) were used for cDNA synthesis and GAPDH was used as internal control. Quantifications were done by Applied Biosystems 7300 Real Time PCR system and fold change calculated using the ΔΔCt method.

### RNA sequencing

Total RNA was extracted using miRNeasy kit (Qiagen) following the manufacturer’s instructions. RNA-Seq libraries were constructed using the TruSeq sample Prep Kit V2 (Illumina). Briefly, 1-2 μg of purified RNA was poly-A selected and fragmented with fragmentation enzyme. After first and second strand synthesis from a template of poly-A selected/fragmented RNA, other procedures from end-repair to PCR amplification were done according to library construction steps. Libraries were purified and validated for appropriate size on a 2100 Bioanalyzer High Sensitivity DNA chip (Agilent Technologies.) The DNA library was quantified using Qubit and normalized to 4 nM before pooling. Libraries were pooled in an equimolar fashion and diluted to 10 pM. Library pools were clustered and run on Nextseq500 platform with paired end reads of 75 bases, according to the manufacturer’s recommended protocol (Illumina). Raw reads passing the Illumina RTA quality filter were pre-processed using FASTQC for sequencing base quality control. Sequence reads were mapped to UCSC human genome build using TopHat and differential gene expression determined using Cufflinks 2.1.1 and Cuffdiff2.1.1 as implemented in Base- Space.

### Lentiviral Transduction

Lentiviral particles were generated by transfecting HEK-293T cells with 8μg of expression vectors (MIR200 CRISPR V2, miR-Zip control, miR-Zip-200c) and 2μg each of packaging vectors (pMDL, pVsVg and pRevRes) in complex with 24μg PEI (Polysciences) in 0.9% NaCl. Viral titer was allowed to concentrate for 48 hours, after which supernatant was collected, centrifuged at 400g for 5 minutes and passed through 0.22μm syringe filter. For transduction, cells were seeded in 6-well plate (100,000 to 150,000) and infected in the presence of 8μg/ml polybrene (Sigma). Cells were selected in medium containing 3μg/ml puromycin (Sigma) and cultured in 1 μg/ml puromycin. MIR200 CRISPR V2 was custom-made and purchased from Genscript (sequence of the gRNA is TGGGAGTCTCTAATACTGCC). miR-Zip control and miR-Zip-200c plasmids were purchased from System Biosciences.

### CRISPR/Cas9 library lentiviral generation and viral transduction

Lentivirus was generated with HEK293TN cells in T175 flasks. Briefly, 9.2μg pMDLg/pRRE plasmid (Addgene, #12251), 4.6μg pRSV-REV plasmid (Addgene #12253), 4.6μg pMD2.G plasmid (Addgene #12259) and 13.8μg of Human CRISPR Knockout Pooled Library (GeCKO v2) (Addgene # 1000000049) part A or part B were combined with Lipofectamine 3000 (Thermo Fisher Scientific # L3000015) according to the manufacturer’s protocol, and transfected the cells with the mixture. The virus medium was collected 24h after transfection and centrifuged at 2000 rmp for 10 minutes to remove cells and debris. Then the supernatant was filtered with a 0.45μM pore filter and frozen at - 80°C. HCT116-Cas9 cells was transduced with serial dilutions of a virus to find the MOI of ~0.3.

### CRISPR screen

HCT116-Cas9 cells were transduced with lentiviral Human GeCKO v2 library part A and part B at MOI of 0.3 in the presence of 4 μg/mL of polybrene for 24h, then replaced the virus medium with fresh growth medium and continued to culture the cells for 48h. The cells were selected with 4 μg/mL of puromycin for 3 days. Then combined both half libraries cells into together and collected 2.5*10^7^ cells for genomic DNA isolation. Next generation sequencing was performed on the Illumina HiSeq 2500 platform in Deep Sequencing Facility of TU Dresden. The raw FASTQ files were analysed with MAGeCK-VISPR.

### Genomic DNA isolation and PCR amplification

Genomic DNA was extracted with NucleoSpin® Blood XL (Machery Nagel # 740950.50) according to the manufacturer’s protocol. The first round PCR of Next Generation Sequence (NGS) is performed with 26 separate 100-μL redundant reactions, each containing 5μg of DNA, 50μL Q5® Hot Start High-Fidelity 2X Master Mix (NEB # M0494L), and 5μL of a 10μM solution of each primer (P5 and P7). The PCR amplification program was as follows: step 1, 98°C for 30s; step 2, 98°C for 10s; step 3, 62°C for 30s; and step 4, 72°C for 30s; steps 2-4 is repeated for 25 times, step 5, 72°C for 2min. Then the PCR production was sent to NGS.

Primer P5: ACACTCTTTCCCTACACGACGCTCTTCCGATCTNNNNNTCTTGTGGAAAGGACGAAACACCG

Primer P7: GTGACTGGAGTTCAGACGTGTGCTCTTCCGATCTTCTACTATTCTTTCCCCTGCACTGT

### Survival analysis

Gene expression profiles of colorectal cancer patient samples were obtained as normalized values from GEO (GSE39582 and GSE33113) and mRNA z-score values for TCGA profile (TCGA, Nature 2012) from cbioportal platform. MS score in patient samples was calculated as a difference of averaged z-score of up-regulated and down-regulated genes identified from RNA sequencing of FACS sorted high and low population of miR-200b/c sensor expressing cells. Samples were then categorized as MS-low and MS-high based on the median value to generate survival curve using Kaplan-Meier estimate. Significance between the two groups was assessed using log-rank test in R software.

### Gene Set Enrichment Analysis

Gene set enrichment analysis (GSEA) was performed using GSEA 4.0.3 software for the association of miR sensor up-regulated genes with ‘Hallmark EMT gene set’ in a patient gene expression profile obtained from GEO database (GSE39852 and GSE33113) and cbioportal platform for TCGA profile. miR sensor up regulated genes score was provided as continuous label of averaged z-score values and calculated the gene ranks using Pearson ranked gene metric.

### Gene signature activation and associational analysis

EMT, TNF-α, G2M and E2F related gene signatures were obtained from MSigDb v7.1 and assessed their gene signature activity using z-score method in colorectal patient mRNA expression profiles (GSE33113 and GSE41258). When up- and down-regulated gene signatures information are available for the study, a difference of z-score between up- and down-regulated gene signatures was carried out. With the z-score activity of the signatures across the colorectal samples, correlation analysis was performed in R using cor.mat function to assess the significance of association and plotted the graph using GraphPad Prism 8. For the sensitivity analysis of miR-sensor RNA seq genes, the difference between up- and down-regulated genes z-score activity was used to categorize the samples into low and high patient sample groups based on the median value in the colorectal cancer patient expression profile (GSE81980). miR-200c expression for the matched mRNA expression profile was obtained from GSE81981 to categorize the samples into low and high based on the median expression for the enrichment of Hallmark genesets.

## RESULTS

### The sensor plasmid can detect endogenous miR-200b/c activity

We tested the ability of the dual fluorescence sensor to accurately report endogenous levels of miR-200b/c when overexpressed by transient transfection. As a cellular model, we chose the colorectal cancer cell line HCT116, which expresses high levels of miR-200s (17) and has an epithelial-like phenotype (29). HCT116 cells were co-transfected with R^mut^G^mut^ control plasmid in the presence of miR-200c mimics (pre-miR-200c) or scrambled control (pre-Control), and the green and red fluorescence intensity were recorded on a flow cytometer. As a result, we found that the double mutant plasmid allowed a robust and coordinated expression of the fluorescent transgenes (cells along a diagonal line in the FACS plot), an important prerequisite for the detection of changes of signal intensities due to the binding of endogenous miRNAs. Overexpression of pre-miR-200c produced no detectable alteration of fluorescence in these cells, **Suppl. Figure 1A**, indicating the lack of binding activity. By contrast, using the signal from the non-binding R^mut^G^mut^ plasmid to set the FACS gates, we could observe the appearance of a population with reduced red intensity in cells transfected with the R^wt^G^mut^ sensor (with scrambled control), **Figure 1C**. This suggests that, when expressed in cells with detectable levels, the miR-200b/c sensor can report a significant binding activity in a sub-population of cells, potentially carrying the highest miR-200b/c levels. Similar results were obtained from another miR-200 positive cell line, the human bladder carcinoma RT112, see **Suppl. Figure 1B**. In HCT116 cells, the percentage of cells in the gated area for high miR-200b/c further increased upon miR-200c overexpression, **Figure 1C**, suggesting the notion that the sensor can be used to monitor cells with high miR-200b/c from endogenous and exogenous source.

### The sensor reports miR-200 activity in cells with differential EMT states

The miR-200 family have a strong EMT-repressing role and are normally reduced in mesenchymal-like cells (18). We sought to determine the activity of the sensor in cells with altered EMT status using an isogenic cellular system with high and low miR-200. For this, we tested mouse pancreatic cancer cell lines derived from a Pdx1-cre;Kras^LSL.G12D/+^;Tp53^LSL.R172H/+^ mouse model (KPC), which has been further modified to obtain the Zeb1 knockout, called KPCZ, to show a role of Zeb1 in the metastatic cascade of pancreatic tumors (28). The mature sequence of miR-200b/c is highly conserved between human and mice (30). Cells obtained from KPCZ tumors showed marked epithelial-like phenotype and higher E-cadherin level, **Figure 1D** and **Suppl. Figure 2A**, and, compared to mesenchymal-like KPC cells, also increased expression of miR-200 members b and c, **Suppl. Figure 2B**. Upon transfection with miRNA sensors, KPCZ cells showed a superior inhibition rate, see **Suppl. Figure 1C**, and **Figure 1E**, in line with the data obtained with the pre-miR transfection experiment.

To further test the specificity of the detection, we used a CRISPR/Cas9 based approach to permanently knockout miR-200c, the most prominent EMT-suppressing member of the miRNA family (31), taking advantage of a protospacer adjacent motif (PAM) sequence adjacent to the miRNA-200c seed match, **Suppl. Figure 3A**. As a result, MIR200C-KO HCT116 cells showed a drastic reduction of miR-200c levels, as evaluated by qPCR, **Suppl. Figure 3B**. In light of the high sequence homology between the miR-200b and c, we also verified the effect of the KO on the MIR200B gene, and a significant >2-fold reduction was also detected, **Suppl. Figure 3C**. DNA sequencing on single-cell clones obtained from MIR200C-KO cells showed alterations around the PAM in the miR-200c, but not in the miR-200b region, **Suppl. Figure 3D**, suggesting that the observed reduction in miR-200b is likely an indirect effect due to EMT induction. Phenotypic analysis of the MIR200C-KO cells, in fact, showed that they had undergone EMT, as indicated by a pronounced mesenchymal-like morphology and dispersed growth pattern, and confirmed by western blotting and immunofluorescence staining (**Figure 1F** and **Suppl. Figure 4A**). To check that EMT was propelled by the miR-200c knockout and not by an unwanted off-target effect (32), we reconstituted the cells with exogenous miR-200c by transient transfection and could observe an attenuation of the phenotype, **Suppl. Figure 4B**. Once transfected with miR-200b/c sensors, the miR-200C-KO cells displayed a significant reduction in the percentage of cells with inhibition of red fluorescence, **Figure 1G-H**, further supporting the high specificity of the miR-200b/c sensor system. Overall, these data indicated that the sensor can monitor changes in the miR-200b/c levels in living cells with distinct EMT states.

### Utility of the sensor to sort cells by endogenous miR-200b/c levels/EMT status

We then tested the possibility to use the sensor for FACS-sorting cells with a differential EMT status based on the endogenous high and low miR-200b/c expression. HCT116 cells were transfected with R^wt^G^mut^ and R^mut^G^mut^ plasmids on a larger scale and separated by their R^wt^G^mut^ non-inhibited (miR-200b/c low) and red-inhibited (miR-200b/c high) population, **Figure 2A**. Gated areas were selected to collect cells with the same intensity of green fluorescence (the mutant control) and a significant difference in red. After sorting, cells were re-plated and allowed to attach overnight to exclude dead cells from the further characterizations. Initially, a western blot was conducted to monitor the expression of the epithelial marker E-cadherin. No change in E-cadherin abundance was found in R^wt^G^mut^ compared to R^mut^G^mut^ transfected cells (unsorted), to control that the miR-200b/c sensor itself did not alter the EMT phenotype. Analysis of sorted cells showed that E-cadherin levels were unaltered in R^mut^G^mut^ compared to parental cells, while a significant reduction was found in miR-200b/c low, and a relatively marked increase was detected in miR-200b/c high cells obtained from R^wt^G^mut^ transfection, **Figure 2B**. This was the first indication that the sensor-guided FACS sorting strategy was capable of physically separating cells with distinct EMT properties. In a separate experiment, qPCR analysis confirmed the differential gene expression of miR-200c and EMT markers in miR-200b/c high and low sorted cells (**Figure 2C**). Similar results were obtained sorting sensor-transfected RT112 cells, **Suppl. Figure 5A-B**. In addition, we found that sorted high and low cells had a strong propensity for quickly reverting their fluorescence after re-plating, as quantified by FACS analysis and video imaging (**Suppl. Figure 5C-D**), indicating that miRNA activity can be monitored for a few days in vitro in individual sensor-transfected cells.

**Figure 2.**
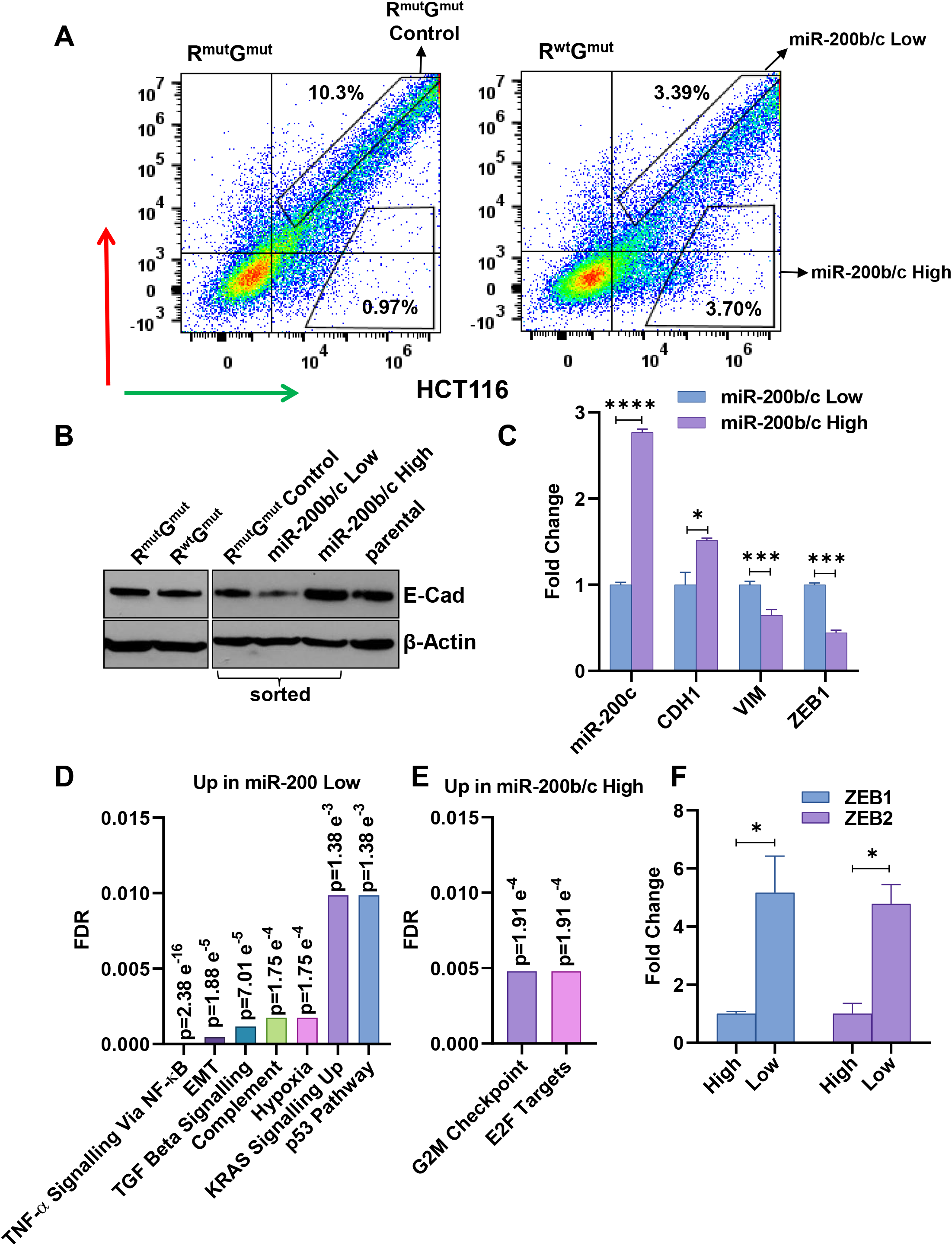
The sensor can be used to sort cells by endogenous miR-200b/c levels. A) FACS plots showing the gates used for sensor-sorting miR-200b/c low and high (right panel) cells after transfection with the R^wt^G^mut^ plasmid. Left panel shows R^mut^G^mut^ transfected control cells and their gate for sorting control. B) Western blot quantification of E-Cadherin in HCT116 cells transfected either with R^mut^G^mut^ or R^wt^G^mut^ plasmids (unsorted, left panel), cells sorted with the gates shown in A) (right panel). C) qPCR quantification of relative mRNA levels of CDH1 (gene coding E-Cadherin), VIM (Vimentin), ZEB1 and miR-200c in miR-200b/c low or high HCT116 cells sorted as in (A). Gene ontology (GO) analysis of RNA sequencing data depicting pathways upregulated in sensor-sorted miR-200b/c D) low and E) high cells. F) qPCR validation of RNA sequencing data. ZEB1 and ZEB2 were quantified using GAPDH as a housekeeping gene. p-values are from students *t*-test. *<0.05, **<0.01, ***<0.001, ****<0.0001

### miR-200b/c sensor-sorted cells can be used to molecularly characterize endogenous EMT states

EMT is a complex multifactorial process. Although a few markers like E-Cadherin, vimentin and ZEB1 can represent a good surrogate for the determination of the EMT state (33) (17), this should be better concluded from broader, and possibly genome-wide characterizations (34). We, therefore, repeated the sorting experiment twice independently (with duplicates) and subjected all the 8 RNAs isolated from high and low cells to sequencing. Using a cut-off of 2-fold for defining differentially expressed genes, we identified 224 down-regulated and 73 up-regulated genes in the 200b/c high compared to low cells, as an overlap between the two distinct sorting experiments, see **Suppl. Figure 6A** and **Suppl. Table 1**. A gene set enrichment analysis revealed that the signatures most significantly enriched in the miR-200b/c low cells were belonging to the TNF-alpha signalling via NF-κB, to EMT, and the TGF-Beta signalling pathways, **Figure 2D**. On the other hand, analysis of the genes upregulated in miR-200b/c high cells indicated the significant prevalence of cell-cycle related targets, like those linked with the E2F transcription factors and genes involved in the G2/M progression through the cells division cycle, **Figure 2E**. Of note, ZEB1 and ZEB2 were identified among the most differentially expressed genes, **Suppl. Figure 6B**. This is particularly important as a further assay validation, since the sensor plasmid is designed as a functional readout of the miR-200b/c binding to the ZEBs transcription factors, Figure 1B, and this result was independently validated by qPCR, **Figure 2F**.

We next sought to validate functionally the role on EMT of the gene signatures obtained from sensor-sorted cells. TGF-Beta is an established master EMT inducer (35). TNF-alpha is less characterized for EMT induction, but has also been associated with EMT promotion in colon cancer cell lines (36). We were able to verify a strongly significant association between TNF-alpha and EMT gene signatures in colorectal cancer patients, **Figure 3A** and **Suppl. Figure 6C**.To validate these findings in our cellular system; we measured NF-κB activity by a luciferase reporter assay in MIR200C-KO cells and found it elevated, **Figure 3B**. Moreover, TNF-alpha treatment-induced EMT markers in parental HCT116 cells, **Suppl. Figure 6D** and significantly reduced miR-200c expression levels, **Figure 3C**. These results confirmed that TNF-alpha and NF-κB pathways can influence EMT and miR-200 in HCT116 cells, as well as the robustness of the data generated with the sensor-sorting strategy. Next, we also aimed to validate the proliferative signature obtained from the miR-200b/c high cells. Supporting evidence came from a few pivotal studies showing that miR-200b/c can sustain colorectal cancer cell (CRC) growth (37,38), and strong inverse correlations was confirmed between G2M and E2F gene signatures and EMT in CRC patients, **Figure 3D-E**. As a functional assay, we measured the growth of the MIR-200C-KO HCT116 cells, using an automated real-time imaging system to monitor cellular confluency. The results indicated that the KO cells grew slower compared to control cells, **Figure 3F**. ShRNA-mediated knockdown of miR-200b/c (using lentiviral miR-Zip technology) was carried out and independently confirmed that HCT116 cells were strongly dependent on miR-200 for their growth, **Figure 3G**. This observation was replicated in the CRC cell line, the COLO 205 (**Suppl. Figure 7A**). We then reasoned that, because of their increased proliferative status, miR-200b/c high cells could have been more sensitive to cell cycle inhibition. We, therefore, subjected HCT116 cells to treatment with a potent inhibitor of a cyclin-dependent kinase, CGP-60474, in the presence or absence of the pre-miR-200c. The results indicated that pre-miR-200c transfected cells had a significantly higher growth reduction with CGP 60474, compared to control cells, **Suppl. Figure 7B-C**. In line with this, CGP-60474 induced a dose-dependent down-regulation of miR-200b/c high cells in sensor transfected HCT116, **Suppl. Figure 7D-E**. Altogether, this functional validation corroborated the notion that the miR-200b/c sensor can represent a valuable tool to separate and characterize cells based on their functional EMT states and can be used to discover biological properties related to miRNA activity.

**Figure 3.**
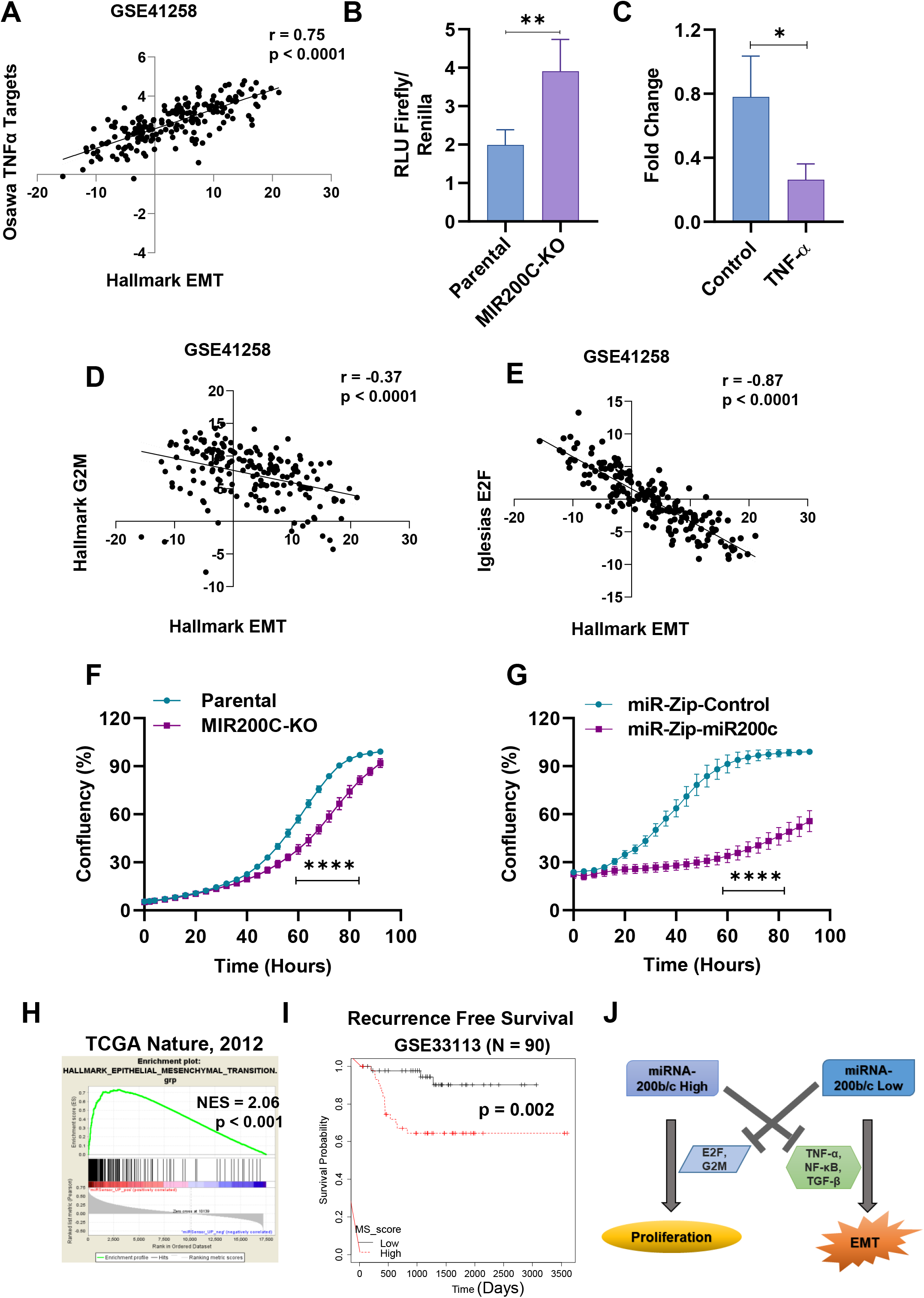
miR-200b/c sensor-sorted cells can be used to molecularly characterize endogenous EMT states. A) Correlation between the z-scores of hallmark EMT and Osawa TNF-α targets in Geo dataset GSE41258 B) Bar graphs showing the relative NF-κB activity (RLU Firefly/Renilla reporter assay) in parental and HCT116 miR-200c KO cells (MIR200C-KO). C) qPCR quantification of miR-200c levels in cells as in B). Cell confluency (proportional to proliferation) measured over time by live cell imaging in D) Correlation between the z-scores of Hallmark EMT and Hallmark G2M targets in GSE41258. E) Correlation between the z-scores of Hallmark EMT and Iglesias E2F targets in GSE41258. Real-time proliferation quantification in F) HCT116 parental and HCT116 MIR200C-KO cells, and G) HCT116 cells stably infected with either miR-Zip control or miR-Zip-200c. H) Gene-set enrichment analysis of miR-200b/c sensor up-regulated genes with hallmark EMT geneset in the indicated dataset as continuous label of miR-200b/c sensor up-regulated scores using Pearson’s gene metric. I) Kaplan-Meier analysis of recurrence free survival in colorectal patients (GSE33113) based on the median value of MS score. J) Scheme of the identified role of miR-200b/c as a molecular switch between EMT and proliferation. p-value was calculated using log-rank test. In B and C the p-values are from students *t*-test. In F and G p-values are from two-way ANOVA and Sidak’s multiple test. *<0.05, **<0.01, ***<0.001, ****<0.0001.

Finally, we performed a gene-set enrichment analysis of miR-200b/c sensor up-regulated genes with hallmark EMT geneset and found very significant association in three independent gene datasets, **Figure 3H** and **Suppl. Figure 8A**, confirming the clinical significance of the data obtained from the sorting experiment. Moreover, when prognostically evaluated, high expression of the gene signature showed to predict poor recurrence-free survival in CRC patients, **Figure 3I** and **Suppl. Figure 8B**. Therefore, the gene signatures obtained from miRNA sensor-sorted cells correlated with clinical EMT and proved to be relevant for prognostic studies. Finally, to further validate the relevance of the method to explore miRNA functions, we tested if the associations identified with the sensor-based approach could have been predicted by an expression analysis, for instance by dividing CRC patients in two groups based on miR-200c expression. As a result, we found that this approach failed to significantly identify TNF-alpha and G2M/E2F signatures as enriched in patients with low and high miR-200c, respectively, **Suppl. Figure 9**.

### The sensor can be used to identify miR-200b/c upstream regulators

As we established that the miRNA sensor functions as an effective fluorescence reporter, we tested the possibility of using it to identify the upstream regulators of miR-200b/c. To do so in an unbiased and genome-wide fashion, we deployed a whole-genome CRISPR-Cas9 library platform and combined it with the sensor/sorting strategy. HCT116 cells were first lentivirally infected to overexpress the Cas9 nuclease and then with the virus obtained from the pooled CRISPR-Cas9 library (39). Around 3×10^8^ HCT116-Cas9-library cells were sensor-transfected and FACS sorted in multiple rounds based on their miR-200b/c levels. MiR-200b/c very high population was expected to be enriched with cells carrying the knockout of genes with the ability to directly/indirectly repress miR-200b/c, **Figure 4A**. After sorting, cells were expanded, their DNA isolated, PCR-amplified and sequenced (all gRNAs are barcoded). As a measure of library representation, a total of 107197 gRNAs were detected in unsorted HCT116 cells expressing the CRISPR library (all gRNAs excluding those with no counts) about 90% of the total, in line with other screens that identified a similar amount of life-essential fitness genes (40). In sorted miR-200b/c high cells, this number dropped to 18817, due to the limited amount of cells that could be FACS-sorted, **Suppl. Figure 10**. By comparing each gRNA abundance between control cells and miR-200b/c high cells, we identified a top-10 list of potential miR-200b/c repressors (**Suppl. Table 2**). The presence of two or more independent gRNAs with a significant enrichment in the miR-200b/c high cells was used as an additional statistical selection criteria (**Figure 4B**). To functionally validate that an actual miR-200b/c repressor has been found, we took a functional approach and independently overexpressed the corresponding ORFs from 5 of the genes in HCT116 cells. Following generation of stable overexpressing cells, we quantified miR-200b and miR-200c expression by qPCR and found that three of them (H6PD, GNPDA1 and ZNF687) significantly reduced levels of both miRNA family members (**Figure 4C and D**). Western blot quantification of EMT markers confirmed for all these genes the regulatory effect on EMT phenotype, **Figure 4E**. Notably, the H6PD gene was also found significantly co-expressed with the EMT markers like ZEB1 and vimentin in a CRC patients’ gene expression dataset, **Suppl. Figure 11**.

**Figure 4.**
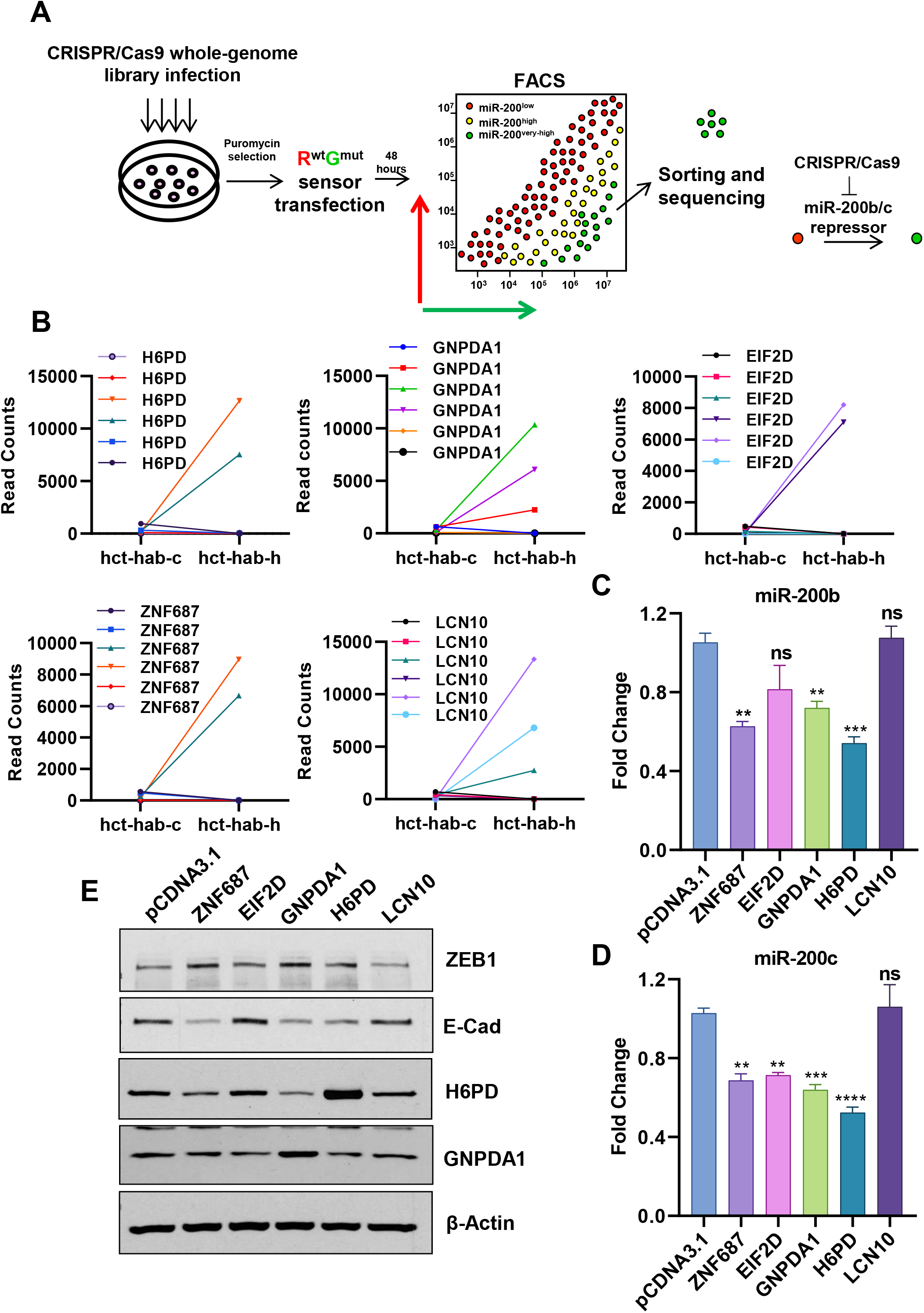
The sensor can be used to identify miR-200b/c upstream regulators. A) Scheme showing the CRISPR-Cas9 screening platform to identify miR-200b/c regulators using the miR-200b/c sensor. B) Graphs showing read counts for the indicated genes’ gRNAs between sorted miR-200b/c high and low cells. C and D) Validation of the top hits from the CRISPR/Cas9 screen by qPCR. miR-200b (C) and miR-200c (D) levels were compared in cells stably transfected with the cDNA clones overexpressing the hits genes, compared to the pCDNA3.1 vector control. E) Western blot quantification of EMT markers ZEB1 and E-cadherin in cells as in C-D). H6PD and GNPDA1 over expression is shown in the respective samples. β-Actin was used as loading control. *<0.05, **<0.01, ***<0.001, ****<0.0001.

Conversely, to test if this technique was also able to identify genes with miR-200b/c-activating functions, we performed an independent screening sorting for the cells with low miR-200b/c (no inhibition of the red fluorescence). Among the top 5 identified (**Suppl. Figure 12A**) we could validate by stable overexpression one gene, HSD17B1, as able to increase the E-cadherin levels in HCT116 cells, and to up-modulate miR-200b and c (**Suppl. Figure 12B-D**).

## DISCUSSION

We report proof-of-concept demonstration that miRNA sensors with natural 3’UTRs and non-binding control sequences can work as functional reporters to detect miRNA expression/activity. They permit the physical separation and the characterization of cells with differential baseline miRNA expression levels, and can be combined with high throughput functional screening technologies of various types. Since miRNAs are key regulators of physiological and pathological mechanisms (41), this system might have important implications in miRNA research and can be optimized to be used in several applications, like miRNA target identification studies or for real-time tracing miRNA activity in single cells in vitro. In the case of miR-200b/c, for instance, the latter feature could be used to visualize the high degree of cellular plasticity normally seen in EMT states (42), as well as in pharmacological studies to screen drugs targeting the EMT process (43).

Nevertheless, the sensor can also be used to infer fundamental biological properties of miRNAs. In terms of ability to distinguish endogenous levels, the RNA profiling results here reported are very encouraging, as the cells with high and low endogenous miR-200b/c matched the corresponding EMT states and the most important elements (like the pro-proliferative role) were independently validated. RNA-sequencing on sensor sorted cells indicated a strong down-regulation of EMT, the TNF-alpha and the TGF-Beta signalling pathways in cells with high miR-200b/c. Both the TNF-alpha pathways and the TGF-Beta signalling have been shown to determine the EMT phenotype of colorectal carcinomas (36,44,45), and can also synergistically converge and cooperate for EMT induction (46), as also revealed by our independent analyses. This implies that the miRNA sensor can be used to study complex molecular associations and identify crucial cell biological mechanisms. In our experimental model, the sensor revealed that miR-200b/c acts as key switch between EMT differentiation and proliferation, being miR-200b/c low cells featured with a strong EMT signature and high miR-200b/c cells more directed towards sustained proliferation, **Figure 3J**. The dualism between EMT and proliferation has been previously suggested or proven in other cancer models (47), and some EMT factors have been found to actively repress proliferation, also in CRC (48,49). miR-200 re-expression is known to induce the colonization via the reverse EMT process (MET) and the promotion of proliferation could facilitate the metastatic out growth (16). miR-200b/c could, therefore, be regarded as a powerful regulator of cellular fate, and deeper investigations on the underlying mechanisms may be crucial in understanding the heterogeneity and the adaptive plasticity frequently observed in tumors (50). To further explore these critical aspects, the miR-200b/c sensor could be in the future combined with single-cell sequencing techniques, contributing to reveal the molecular features of EMT intermediate states (referred to as partial or hybrid EMT), the fundamental importance of which has been recently clearly demonstrated in metastasis (51–53). Of note, the functional associations here identified with the miR-200b/c sensor could not have been predicted with an expression analysis, confirming the importance of activity sensors to investigate miRNA biology.

Another important application is the analysis of the upstream regulation network of miRNAs. MiRNA regulation is primarily investigated by the analysis of specific consensus motifs in the promoter regions for the binding of transcription factors. Our pilot experiments using a whole-genome CRISPR/Cas9 library could identify a few unprecedented miR-200b/c regulators. The low number of hits is likely an effect of the great representation loss due to the sorting of a relatively low number of cells, which could be improved by increasing the sensitivity of the sensor (see below). However, we could functionally validate some genes, which, of note, belonged to ‘unexpected’ pathways, like metabolism. This confirms the power of this unbiased technique and points at a strong connection between EMT and cancer metabolism, which is a very lively field of investigation with several important translational implications (54). H6PD (Hexose-6-phospahate dehydrogenase) is an endoplasmic reticulum (ER) specific enzyme that functions as a glucose 6-phosphate dehydrogenase and converts it to 6-phospho glucano lactone. GNPDA1 (D-glucosamine-6-phosphate deaminase) converts glucosamine-6-phosphate to fructose-6-phosphate and vice versa. These two glucose metabolism enzymes work closely associated (**Suppl. Figure 13A**) and could be further explored in EMT regulation. This is not completely surprising, as the connection between EMT and glucose metabolism has recently started to emerge in the literature (33,55). Remarkably, also the miR-200b/c activator gene here identified, HSD17B1, is a metabolism gene, being part of the testosterone/estrogen synthesizing pathway (**Suppl. Figure 13B**). Estrogen signalling has previously associated with EMT suppression in CRC via the up-regulation of another EMT-repressing miRNA, miR-205 (56), which can work in association with miR-200 (11) and, along the same line, androgen-deprivation therapy has been shown to induce EMT via ZEB1 (57). In conclusion, even if these findings will require further experimental verifications, addressed for instance to understand the direct/indirect nature of the regulation, they highlight the great potentiality of the technique. Future screens performed in larger settings and on improved miRNA sensors (or directly in miRNA reporter cells, as described below) will allow high-resolution functional mapping of the regulatory events upstream of miRNAs.

The miR-200b/c sensor design enables a high level of specificity. The R^mut^G^mut^ and R^wt^G^mut^ plasmids differ only by three-point mutations in each of the three natural miR-200b/c seed matches present in ZEB2 3’UTR, and we previously demonstrated the exclusive binding of the members of the miR-200b/c/429 cluster (27,58). The high level of specificity was here confirmed in the analysis of endogenous levels in different cellular models, and by overexpression and knockdown approaches. However, the sensitivity can be the object of further improvements, allowing to extend the assay to cells with lower miRNA endogenous expression. Sensors designed with artificial 3’UTR with multiple fragments perfect complementary to the targeting miRNA can guarantee a higher level of sensitivity, as previously shown in a modelling study (59). However, the sensor here presented has the advantage of carrying naturally occurring sequences properly controlled with non-binding mutants, in analogy with the dual reporter vectors used to validate bona fide miRNA targets. This method is preferable to better represent physiological conditions (59) by 1) making sure that the sensors are reliably reporting endogenous miRNA activity, i.e. not overestimating their effects, and by 2) minimizing or eliminating miRNA-sponge or decoy effects (21,60), which could alter the biological properties. Another unappreciated factor is that the presence of non-natural 3’UTRs (either containing repeated seed matches or complementary sequences) carrying random spacer sequences increases the chances to introduce novel unwanted, albeit specific, miRNA binding sites. In the absence of non-binding controls, the contribution of these off-target detections can be difficult to estimate. To improve sensitivity, therefore, multiple 3’UTRs controlled with non-binding mutants in a 1:1 ratio could in future be cloned in tandem and tested. Another limiting factor that warrants further improvement is that this sensor can only be transiently introduced in the cells. In fact, after the integration of this vector into the genome it showed recombination events from the highly repetitive sequences. Genome integrated sensors failed to propagate in cells (27), precluding its further use in other applications. To overcome this limitation, future studies should be conducted to improve the sensor plasmids’ architecture, like using single bidirectional promoter vectors (61) to minimize the presence of repetitive sequences. Once improved, this approach could be implemented for all biologically-relevant miRNAs, leading to fundamental discoveries in biomedicine.

## ACCESSION NUMBERS

RNA-sequencing data has been deposited to GEO dataset GSE154429

## ACKNOWLEDGEMENT

We acknowledge the work of Prof. Marcus Peter and Dr. Sun Mi Park for the early development of the sensor plasmids. We would like to thank the FACS-core facility of the Nikolaus-Fiebiger-Zentrum of the Friedrich-Alexander University Erlangen-Nuremberg.

## FUNDING

This work was supported by the Interdisciplinary Center for Clinical Research of the University of Erlangen-Nuremberg (PC) and the German Research Foundation grant DFG-CE-281/5-1 (PC). PC was also supported by IASLC Young Investigator Award. CP and HY were supported by the EU (MSCA grant no.: 861196).

## CONFLICT OF INTEREST

The authors declare no conflict of interest.

---

**Supplementary Fig. 1.**
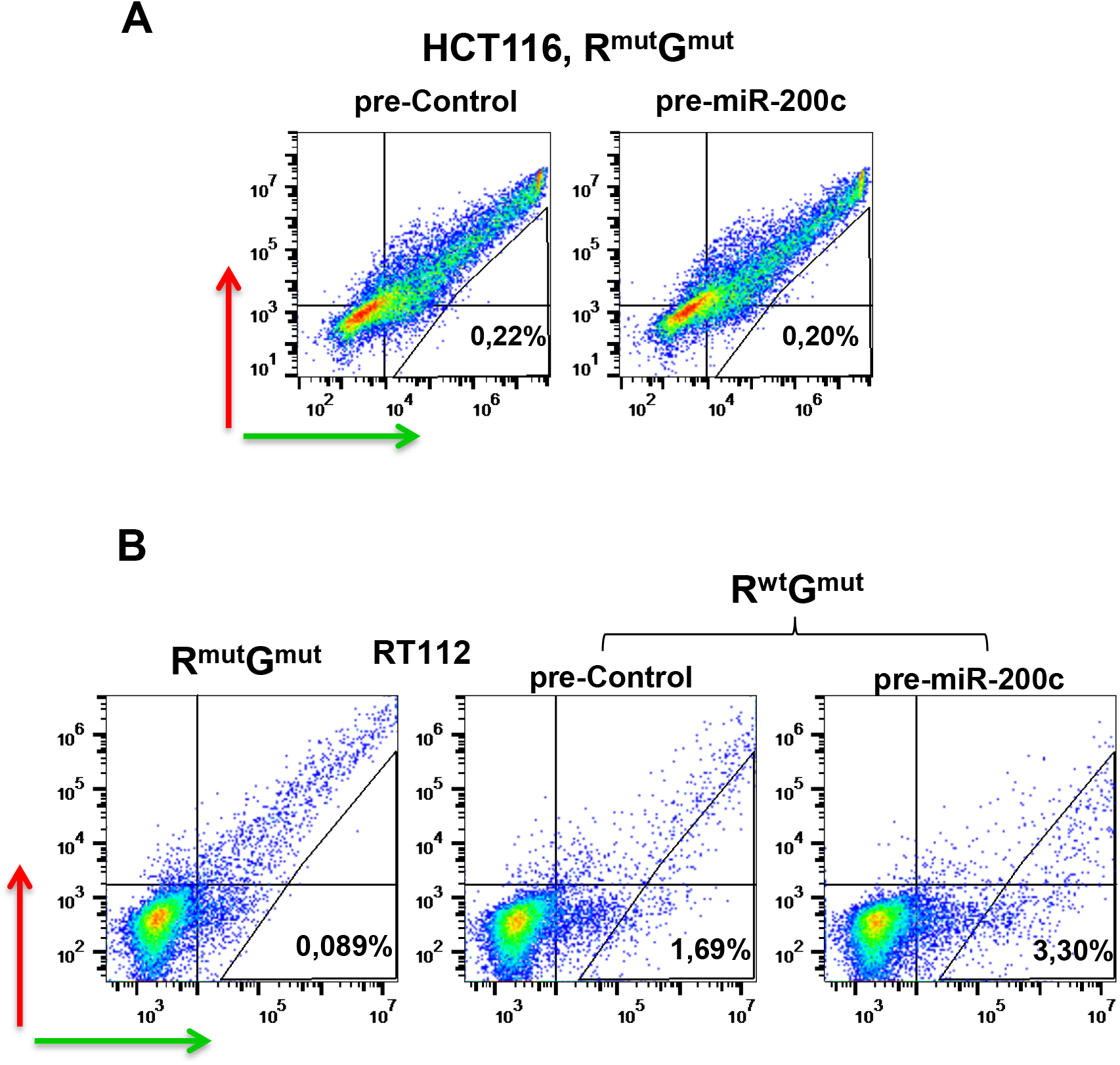
A) FACS plots showing green and red fluorescence in HCT116 transfected with the R^mut^G^mut^ control plasmid in the presence of 100nM pre-control or pre-miR-200c. B) FACS plots showing transfection of RT112 cells with R^mut^G^mut^ or R^wt^G^mut^ plasmids in the presence of pre-control or pre-miR-200c.

**Supplementary Fig. 2.**
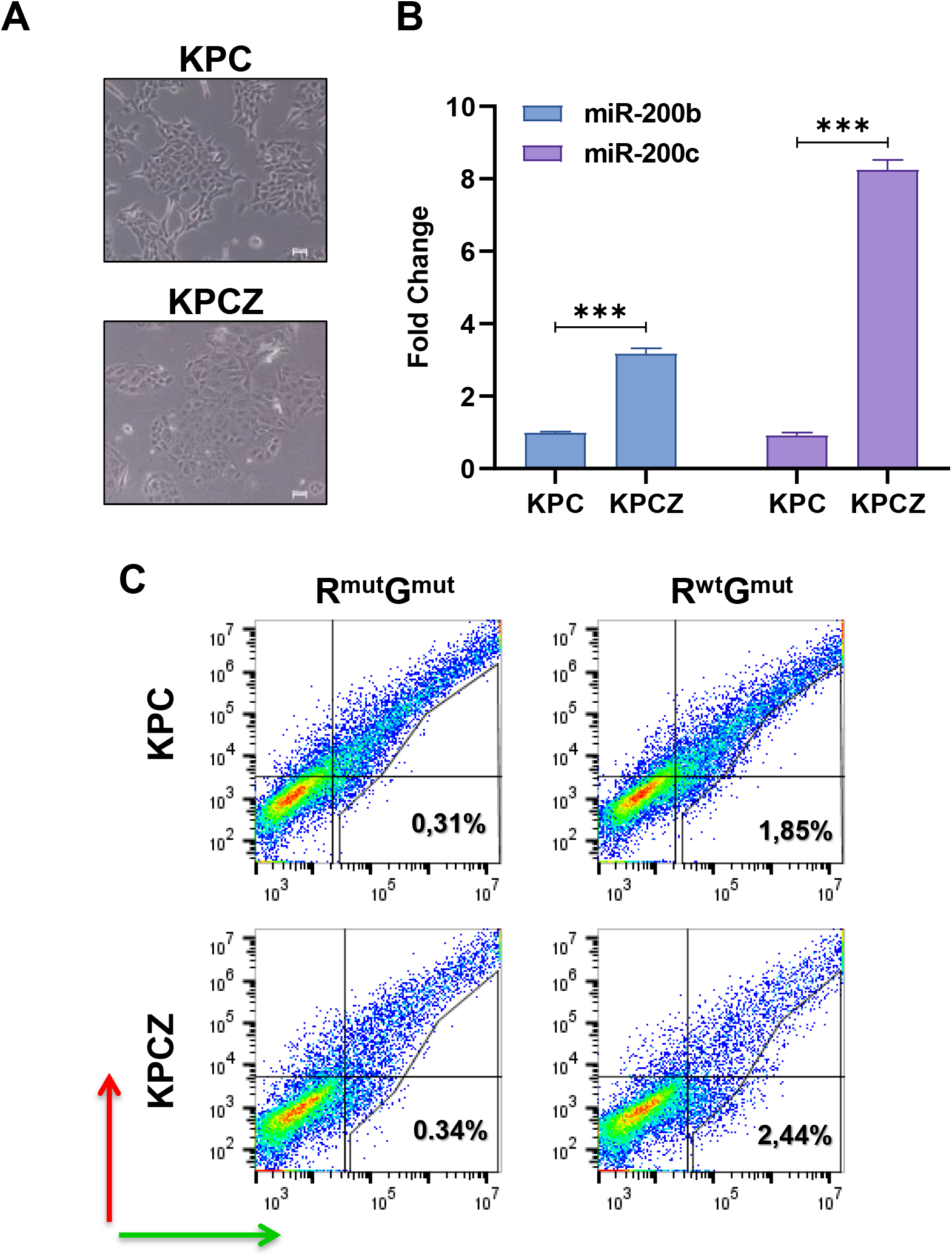
A) Microscopic images of KPC (Pdx1-cre;Kras^LSL.G12D^/+;Tp53^LSL.R172H/+^) and KPCZ (Pdx1-cre;Kras^LSL.G12D^/+;Tp53^LSL.R172H/+^; ZEB^fl/fl^) cells. Scale bar is 50μM. B) qPCR quantification of relative expression levels of miR-200b and miR-200c in KPC and KPCZ cells. C) FACS plots showing transfection of KPC and KPCZ cells with R^mut^G^mut^ or R^wt^G^mut^ plasmids. p-value is from student’s *t*-test.

**Supplementary Fig. 3.**
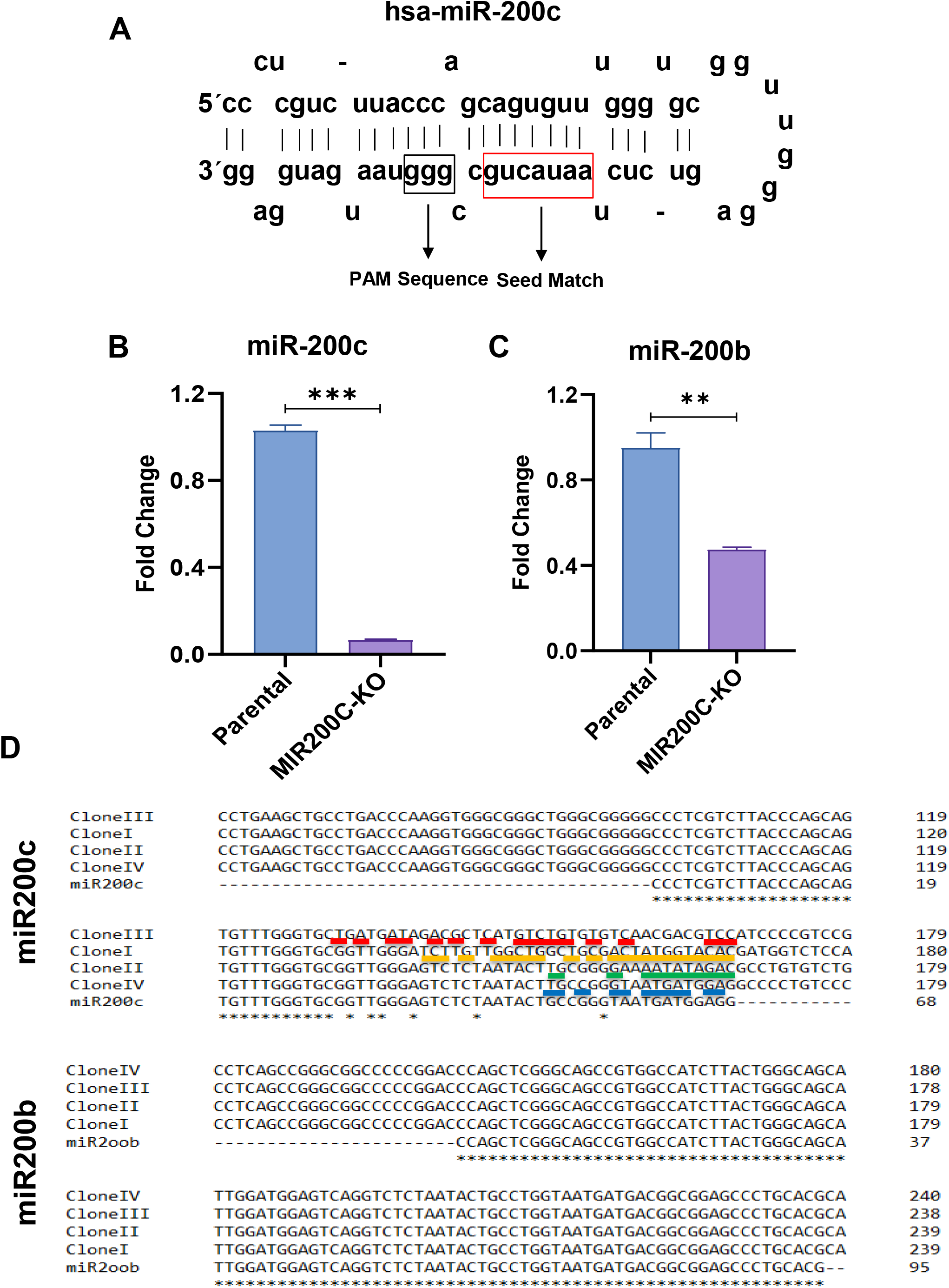
A) Structure of human miR-200c showing the protospacer adjacent motif (PAM) sequence adjacent to the miRNA-200c seed match. qPCR quantification of relative B) miR-200c and C) miR-200b levels upon CRISPR/Cas9 mediated knockout (KO) of miR-200c in HCT116 cells compared to parental cells. p-value is from student’s *t*-test. C) Sequence alignments of single cell clones obtained from HCT116-MIR200C-KO cells comparing with the sequences of miR-200c and miR-200b. Underlined are mutations compared to the reference sequences.

**Supplementary Fig. 4.**
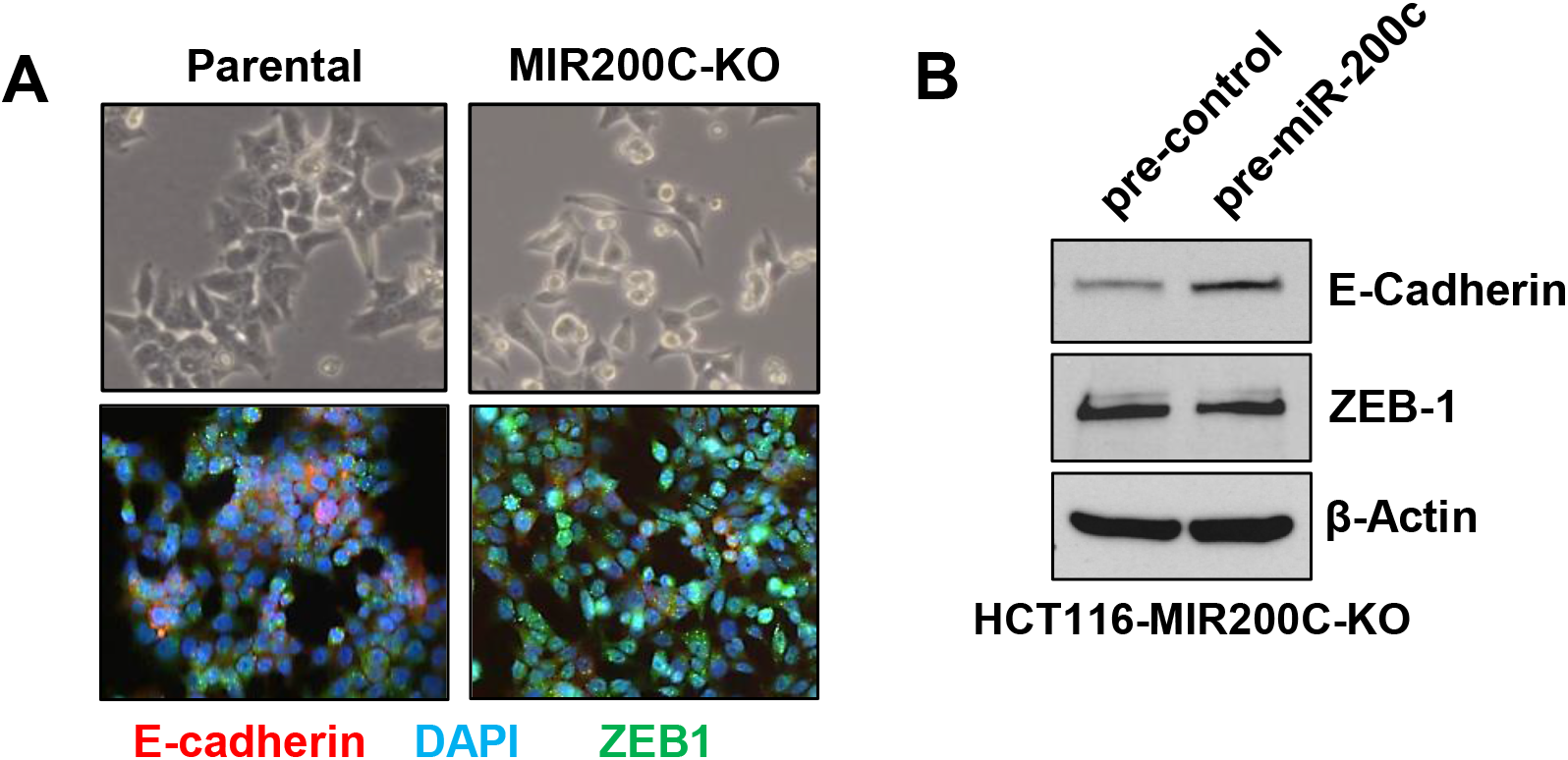
A) Fluorescent microscopic images showing E-cadherin (Red) and ZEB1 (Green) levels in HCT116 parental and MIR200C-KO HCT116 cells. Dapi (Blue) is used to stain the nucleus. B) Western blot quantification of E-cadherin and ZEB1 protein levels in HCT116 MIR200C-KO cells transfected either with pre-miR-control or pre-mir-200c. β-Actin was used as loading control.

**Supplementary Fig. 5.**
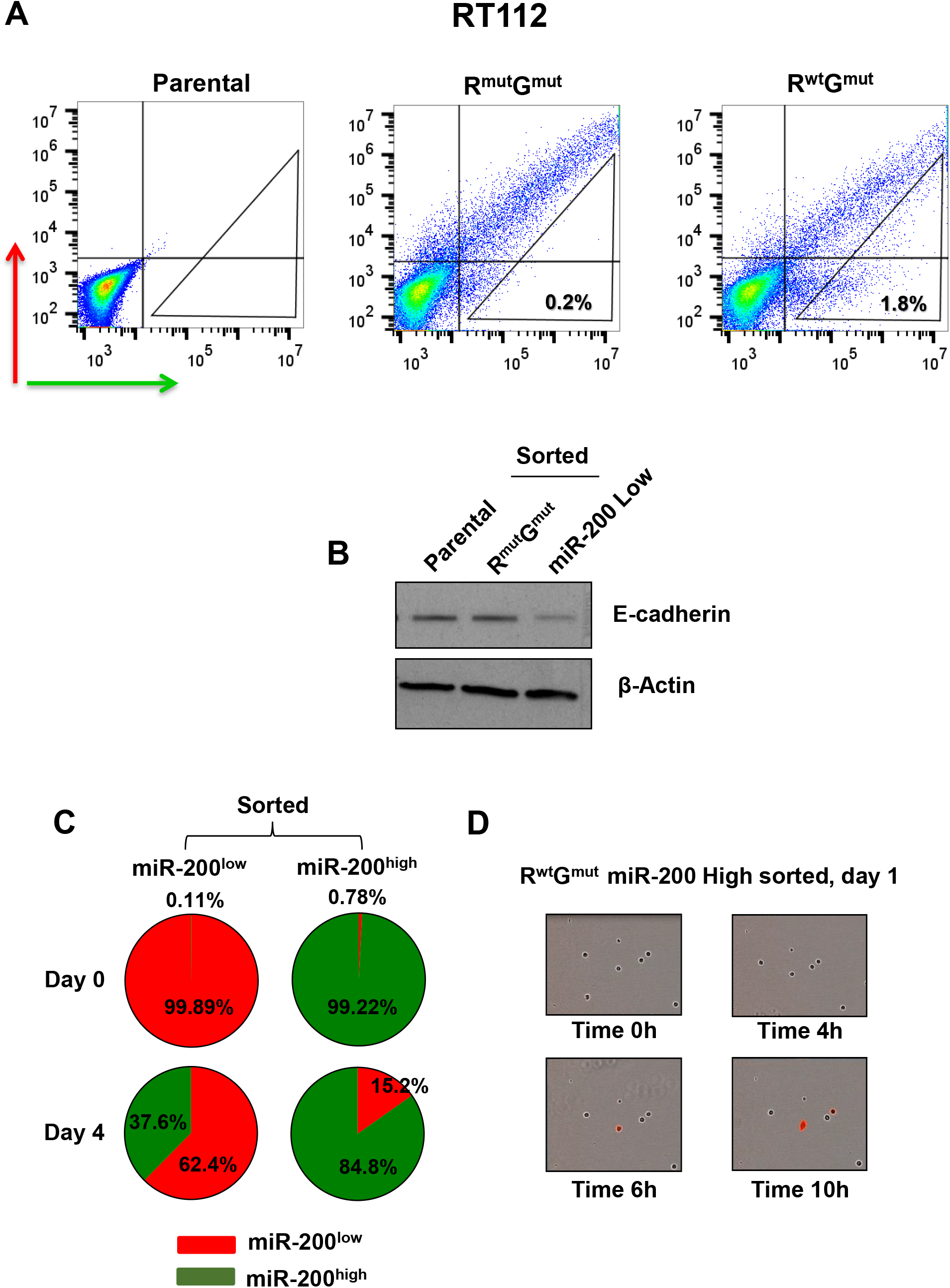
A) FACS plots showing the separation of RT112 cells based on their miR-200b/c levels by using the sensor. B) Western blots showing the levels of E-cadherin and β-Actin (loading control) proteins in parental RT112 cells and in sensor sorted cells. C) Pie charts showing the % of cells with miR-200b/c low and high on day 0 and day 4 of sorting, as evaluated by FACS analysis. D) Images taken with live cell fluorescence imager, showing the sorted miR-200b/c high HCT116 cells increasing red fluorescence.

**Supplementary Fig. 6.**
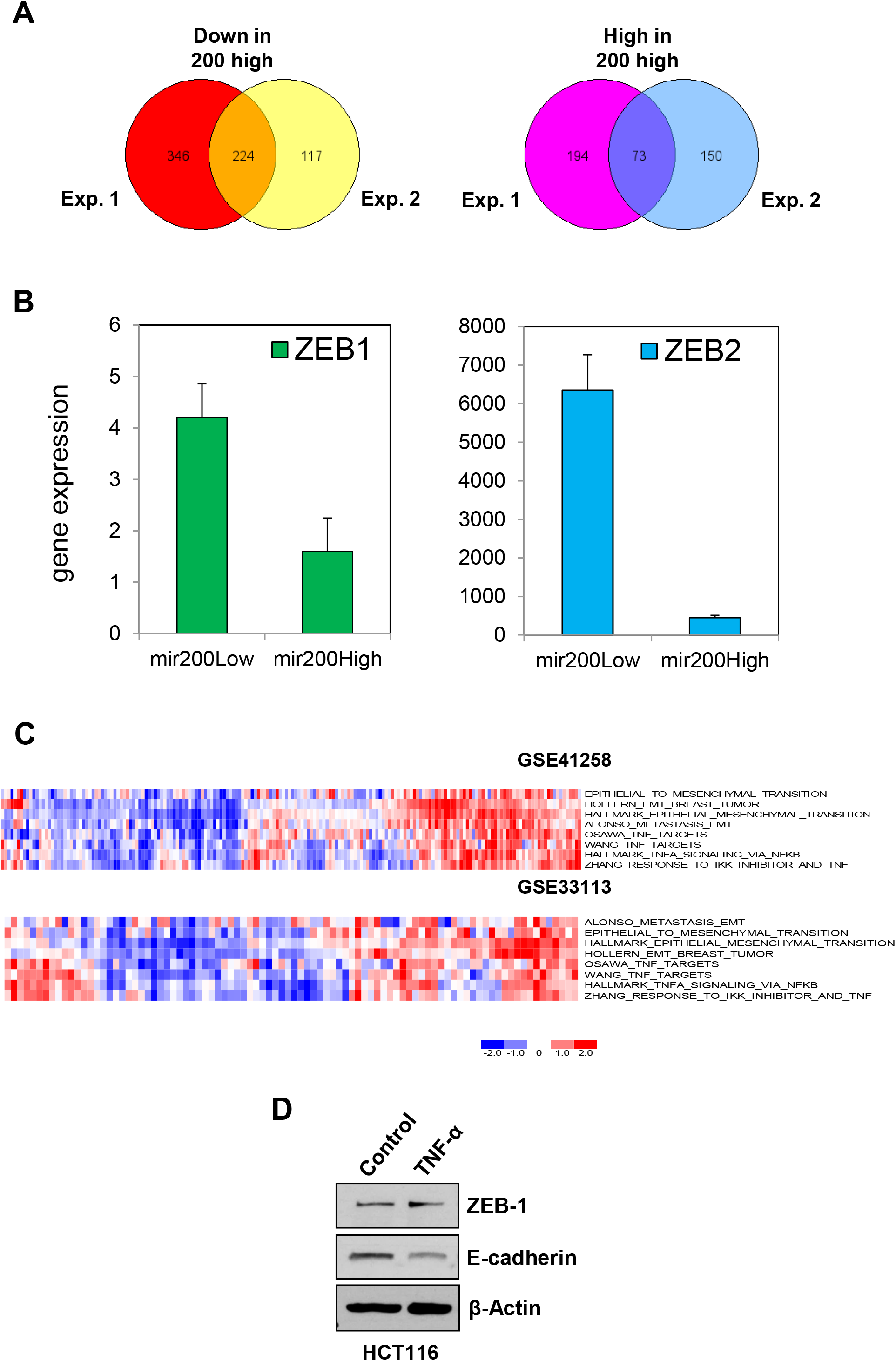
A) Venn-diagram showing overlap between the gene lists obtained from the two independent RNA-sequencing of miR-200b/c high and low cells from the two sorting experiments (Exp.1 and 2). B) ZEB1 and ZEB2 relative expression levels from RNA sequencing obtained from the sensor sorted miR-200b/c high and low cells. C) Heatmap visualization of EMT and TNF-alpha gene signature activity pattern in colorectal cancer samples. Shown are the z-score activity pattern of EMT-associated and TNF-alpha signatures in colorectal cancer gene expression profiles, GSE33113 (N=90) and GSE41258 (N=186), indicating co-regulated activity, ranging from a minimum correlation of 0.35 to a maximum of 0.75 with an adjusted p-value < 0.01. D) Western blot quantification of ZEB1 and E-Cadherin protein levels with or without TNF-alpha treatment (30ng/ml) in HCT116 cells.

**Supplementary Fig. 7.**
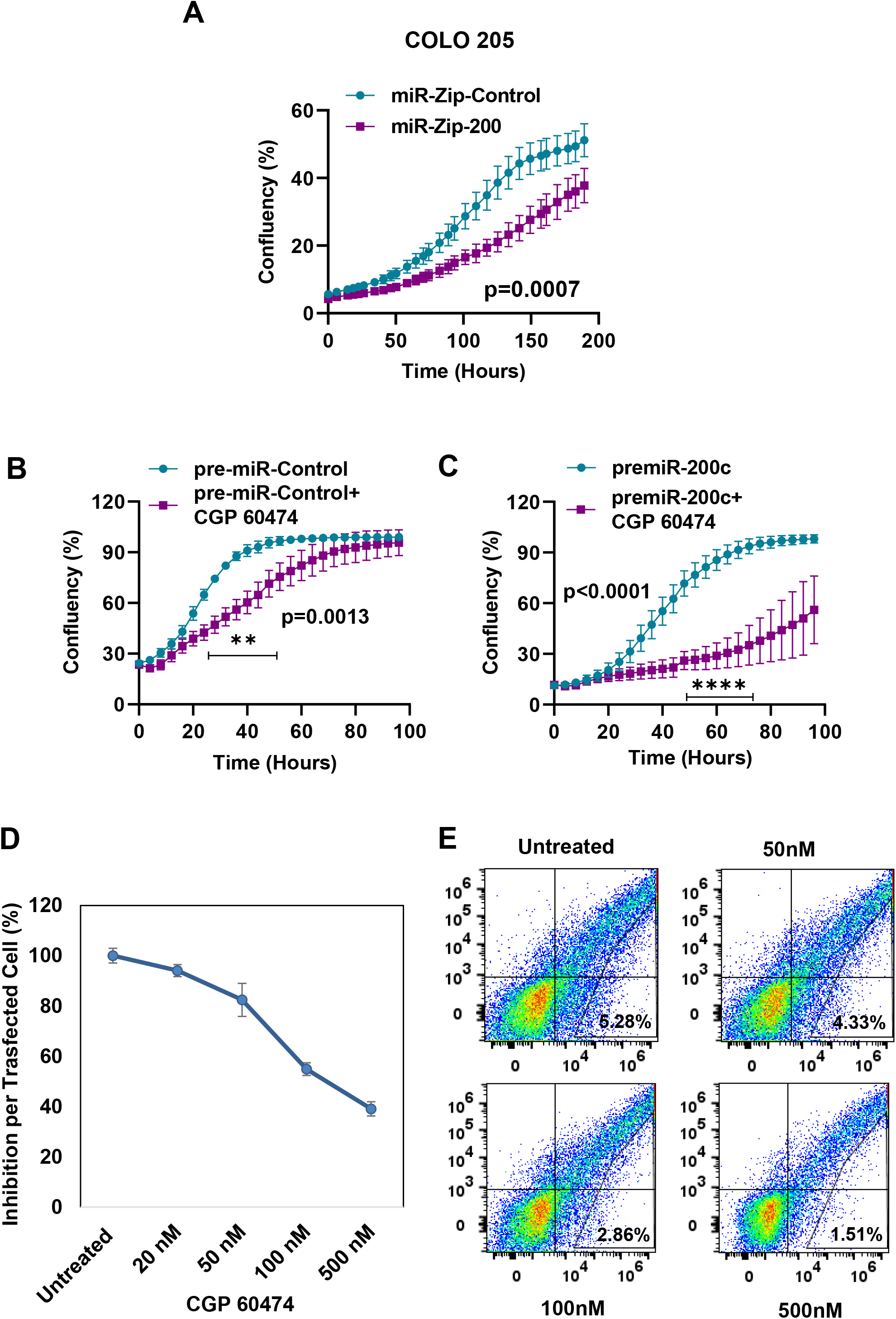
A) Graph showing the proliferation of Colo-205 cells in the presence of miR-Zip control or miR-Zip-200. Cell proliferation in HCT116 cells treated with B) pre-miR-control alone or pre-miR-control+cdk inhibitor (CPG-60474, 20nM) and C) pre-miR-200c alone or pre-miR-200c+cdk inhibitor (20nM). D) Graph showing the % of inhibition in HCT116 cells transfected with the R^wt^G^mut^ plasmid in the presence of the indicated concentrations of Cdk inhibitor CGP-60474. E) FACS plots showing the % of inhibition in HCT116 cells in the presence or absence of increasing doses of CGP-60474. In A, B,C the p-values are from two-way ANOVA and Sidak’s multiple test. *<0.05, **<0.01, ***<0.001, ****<0.0001.

**Supplementary Fig. 8.**
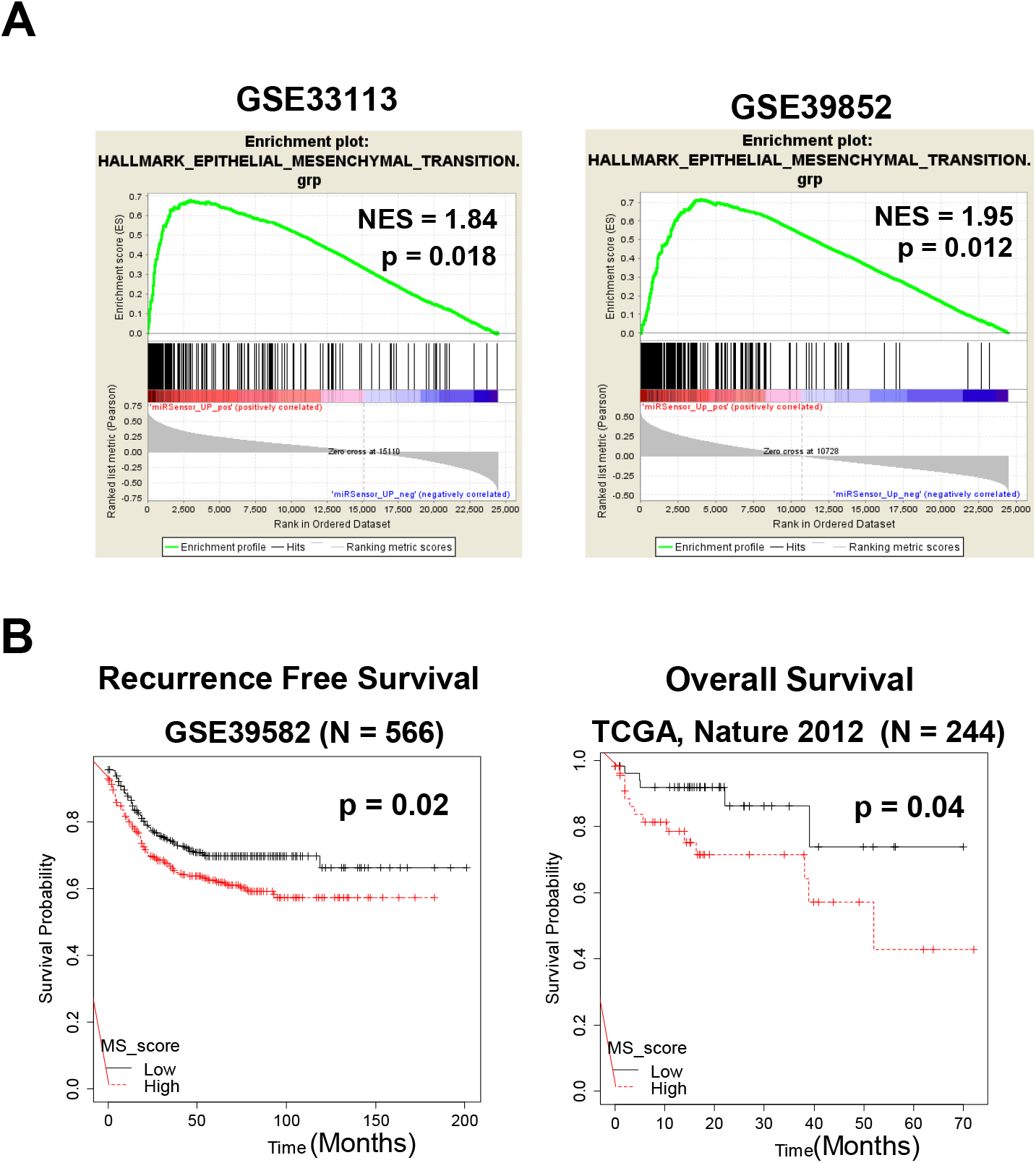
A) Gene-set enrichment analysis of miR-200b/c sensor up-regulated genes with hallmark EMT geneset in the indicated datasets as continuous label of miR-200b/c sensor up-regulated scores using Pearson’s gene metric. B) Kaplan-Meier analysis of recurrence free survival (GSE39582) and overall survival (TCGA, Nature 2012) in colorectal patients based on the median value of MS score. p-values were calculated using log-rank test.

**Supplementary Fig. 9.**
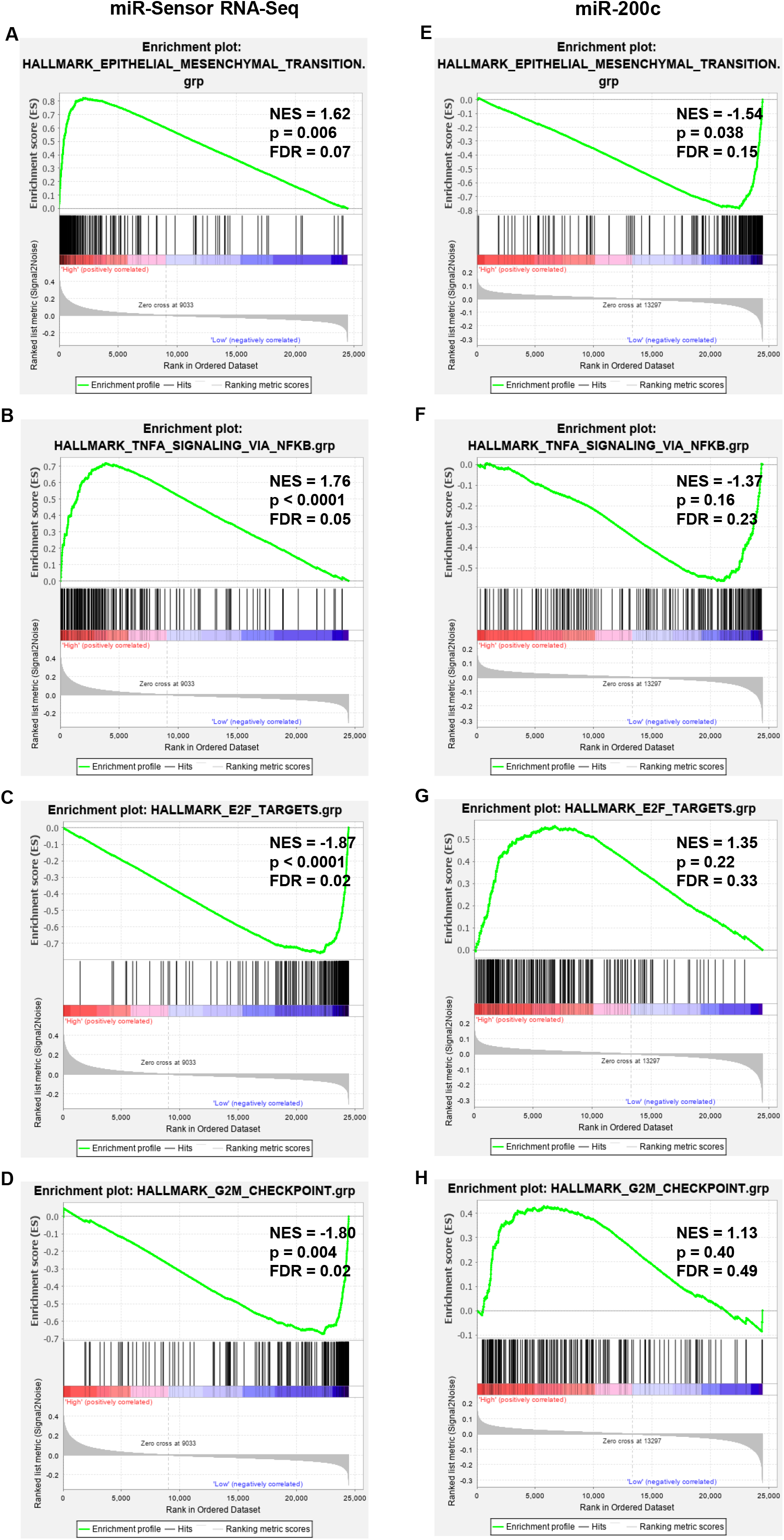
Gene-set enrichment analysis of hallmark genesets (EMT, G2M, E2F and TNFA) with the low and high categorized patient samples based on (A-D) RNA-seq genes obtained from the sensor-sorting experiment, and based on (E-F) miR-200c expression in the GSE81980 dataset. GSEA was performed with the ranking of genes with signal2noise metric.

**Supplementary Fig. 10.**
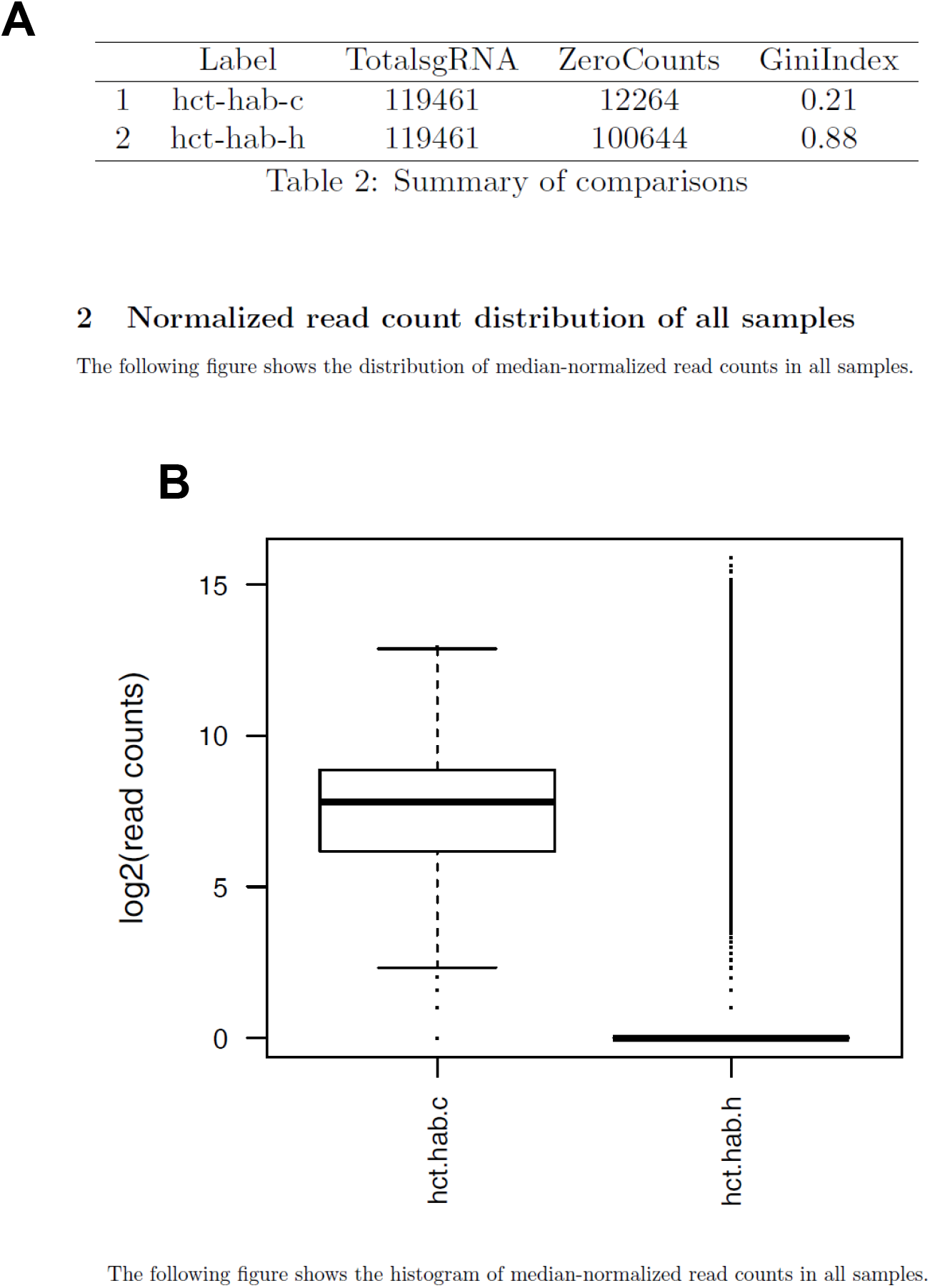
A) Table showing the total gRNAs, the number of absent gRNA (zero counts) in control (hct-hab-c) and miR200b/c-high cells (hct-hab-h). B) Box-plot showing the distribution of read counts in control (hct-hab-c) and miR200b/c-high cells.

**Supplementary Fig. 11.**
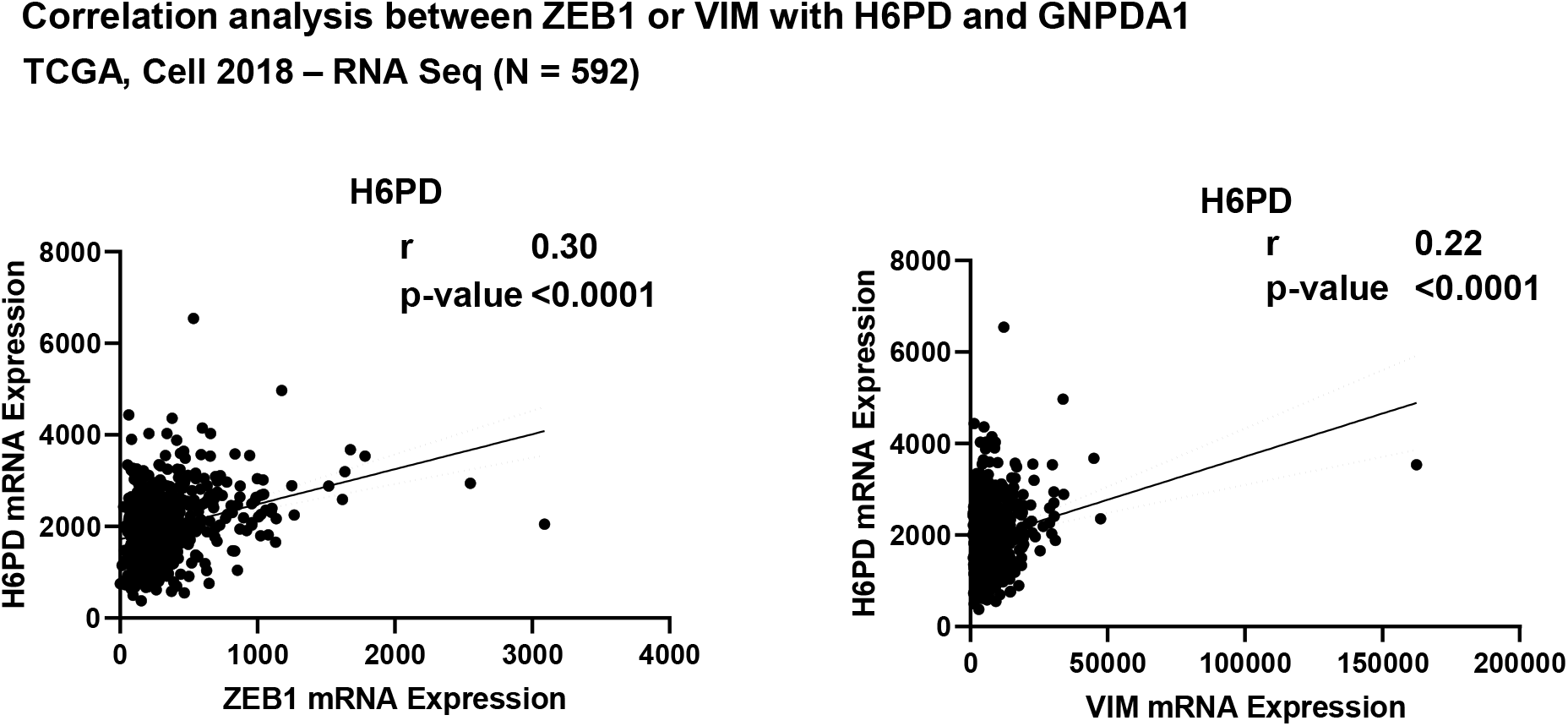
A) Correlation analysis of H6PD mRNA with that of either ZEB1 or Vimentin, in TCGA RNA sequencing data of colorectal adenocarcinoma patients (n=592).

**Supplementary Fig. 12.**
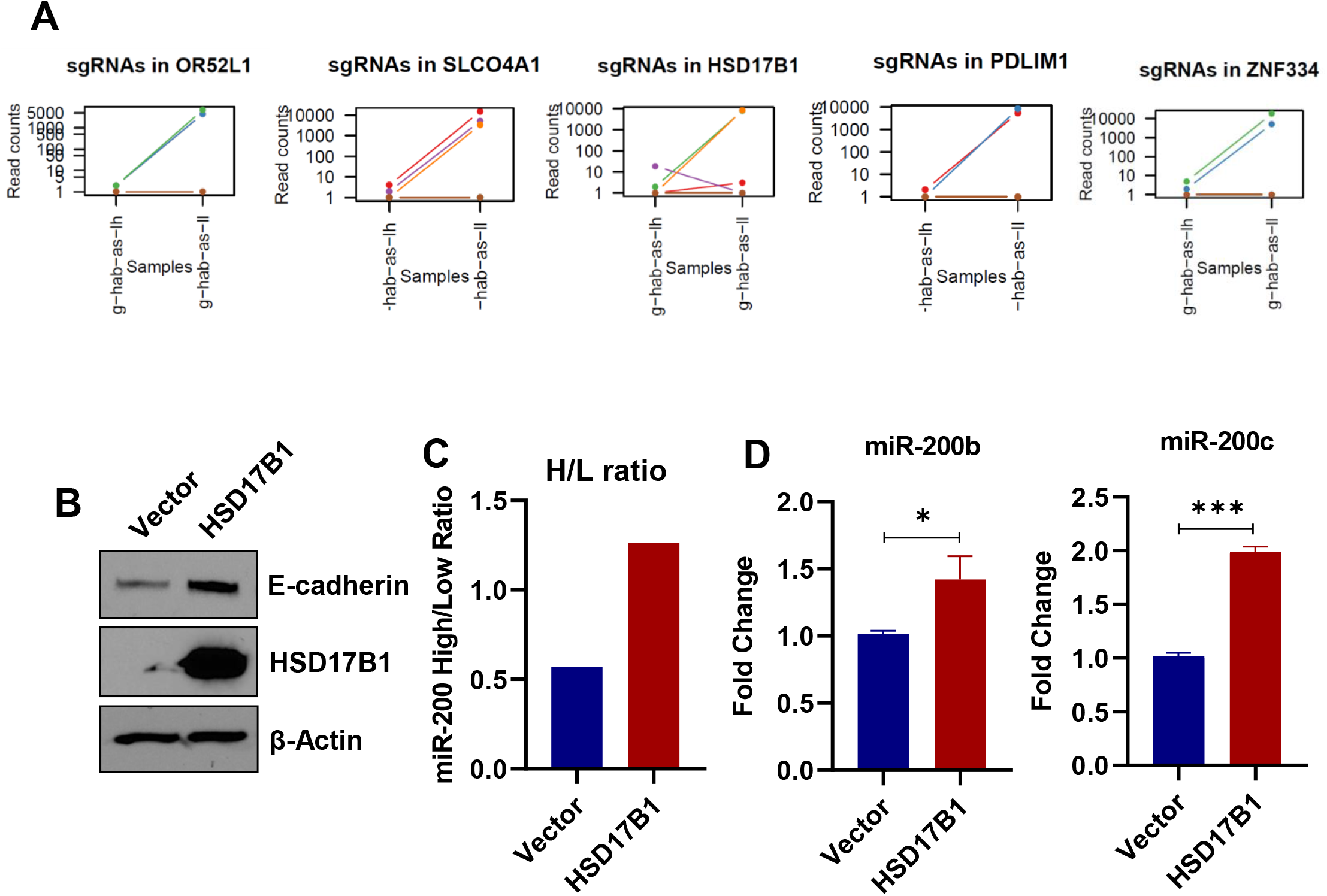
A) Graphs showing the read counts of the gRNAs corresponding to the indicated genes in sensor-sorted miR-200b/c low cells compared to control cells. B) Western quantification of E-cadherin and HSD17B1 levels in HCT116 cells stably transfected with either vector (pCDNA3.1) or HSD17B1 cDNA plasmid. C) Graph showing the ratio of miRNA-200b/c high and low cells in cells as in (B) transfected with the R^wt^G^mut^ plasmid. D) qPCR analysis of miR-200b (left) and miR-200c (right) in HCT116 cells stably transfected with either vector (pCDNA3.1) or HSD17B1 cDNA plasmid. p-values are from students *t*-test.

**Supplementary Fig. 13.**
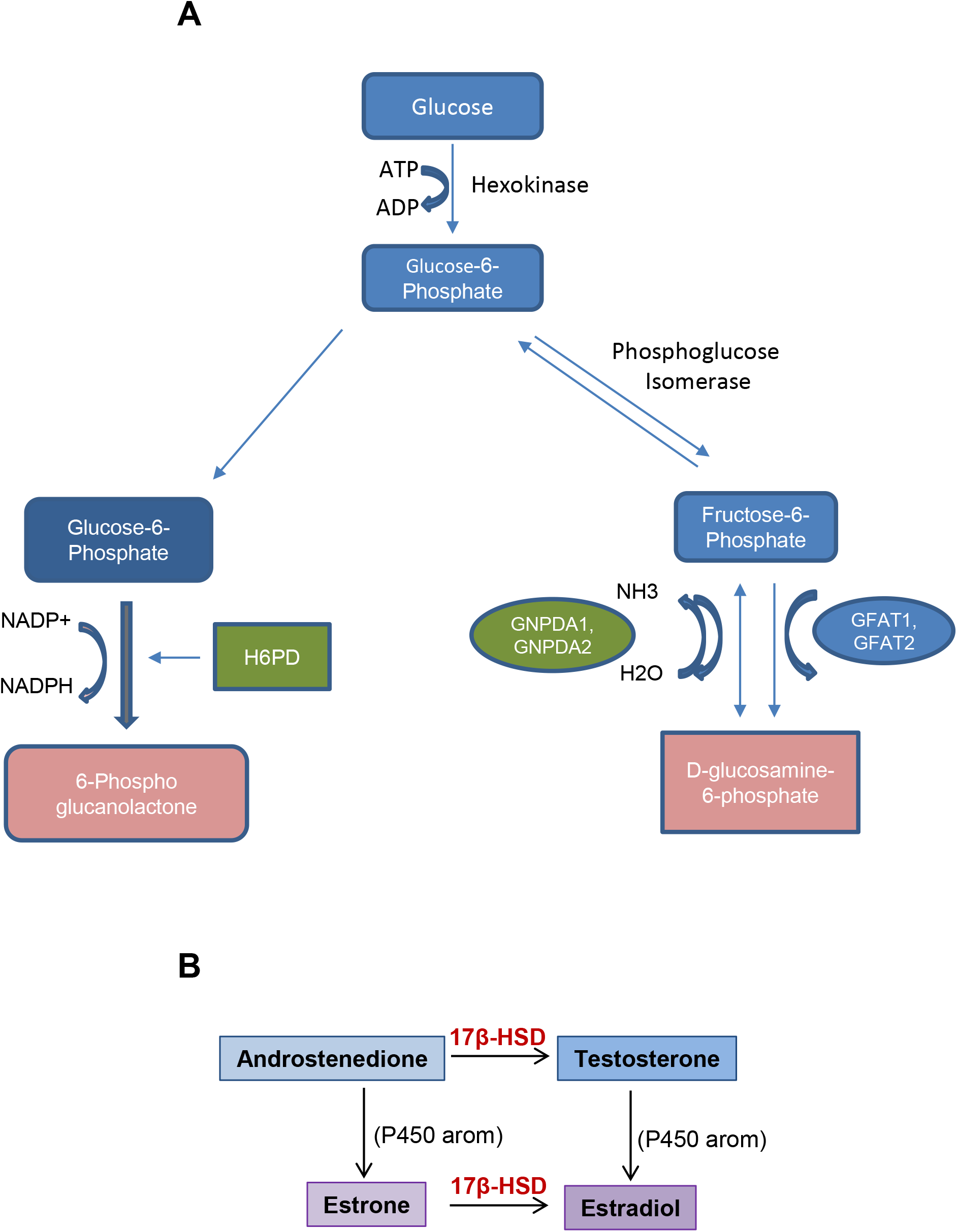
Scheme showing the metabolic functions of A) H6PD and GNPDA1 and B) HSD17B1.

**Supplementary Table 1.**
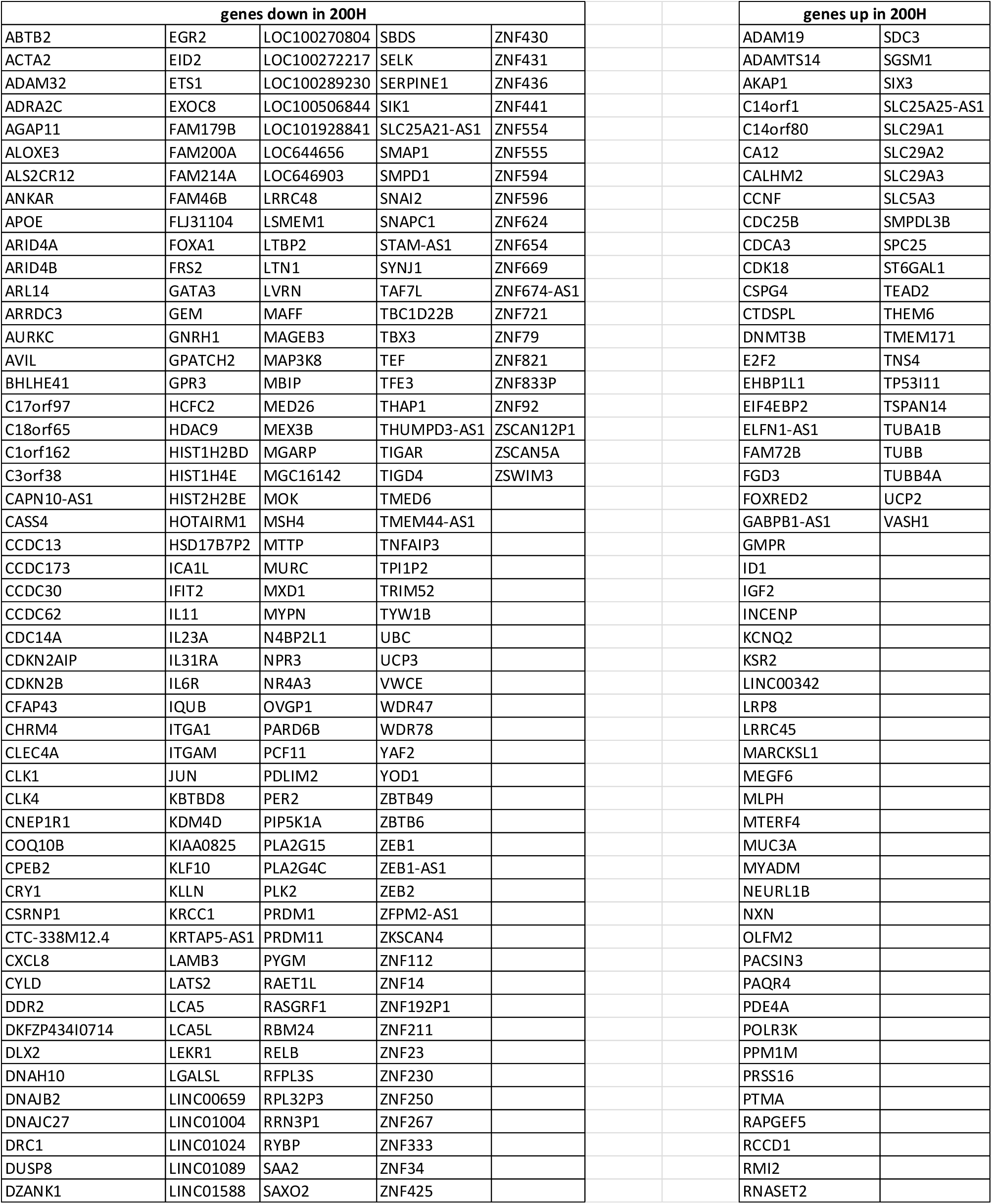
Genes up and down regulated in HCT116 cells sorted for high and low miR-200b/c levels, and subjected to RNA sequencing.

**Supplementary Table 2.** Top 10 hits genes identified by the CRISPR/Cas9 screens of high and low miR-200b/c HCT116 cells.

| top 10 in miR200H | top 10 in miR200L |
| --- | --- |
| ZNF687 | OR52L1 |
| EIF2D | MED10 |
| GNPDA1 | HTN1 |
| H6PD | SLCO4A1 |
| LCN10 | HSD17B1 |
| BTBD9 | SAMD13 |
| CHRNA2 | PDLIM1 |
| F9 | NPC1L1 |
| TRIB3 | TMEM81 |
| PROS1 | ZNF334 |

## Notes

### Competing Interest Statement

The authors have declared no competing interest.

